# Chemoproteomics-Enabled Ligand Screening Yields Covalent RNF114-Based Degraders that Mimic Natural Product Function

**DOI:** 10.1101/2020.07.12.198150

**Authors:** Mai Luo, Jessica N. Spradlin, Scott M. Brittain, Jeffery M. McKenna, John A. Tallarico, Markus Schirle, Thomas J. Maimone, Daniel K. Nomura

**Affiliations:** Department of Chemistry, University of California, Berkeley, Berkeley, CA 94720 USA; Novartis-Berkeley Center for Proteomics and Chemistry Technologies; Novartis Institutes for BioMedical Research, Cambridge, MA 02139 USA; Department of Molecular and Cell Biology, University of California, Berkeley, Berkeley, CA 94720 USA; Department of Nutritional Sciences and Toxicology, University of California, Berkeley, Berkeley, CA 94720 USA; Innovative Genomics Institute, Berkeley, CA 94720 USA

**Keywords:** RNF114, PROTAC, targeted protein degradation, covalent ligand, cysteine, chemoproteomics

## Abstract

The translation of natural product function to fully synthetic small molecules has remained an important process in medicinal chemistry for decades resulting in numerous FDA-approved medicines. We recently discovered that the terpene natural product nimbolide can be utilized as a covalent recruiter of the E3 ubiquitin ligase RNF114 for use in targeted protein degradation (TPD) – a powerful therapeutic modality within modern day drug discovery. Using activity-based protein profiling-enabled covalent ligand screening approaches, we herein report the discovery of fully synthetic RNF114-based recruiter molecules that can also be exploited for PROTAC applications, and demonstrate their utility in degrading therapeutically relevant targets such as BRD4 and BCR-ABL in cells. The identification of simple and easily manipulated drug-like scaffolds that can mimic the function of a complex natural product is beneficial in further expanding the toolbox of E3 ligase recruiters, an area of great importance in drug discovery and chemical biology.

## Introduction

Natural products have remained a cornerstone of drug discovery research for decades resulting in numerous FDA-approved medicines and tools for biomedical research across a wide range of therapeutic areas (Newman and Cragg, 2016). Historically, a large percentage of natural product-inspired medicines have utilized the natural product as a starting point, wherein the tools of synthetic chemistry are used to fine tune compound properties (i.e a semisynthetic approach). Alternatively, the function of natural products can also serve as motivation for the design of fully synthetic small molecules less constrained by availability, synthetic manipulation limitations, and physicochemical and metabolic liabilities. Modern-day examples of this approach include the translation of the alkaloid cytisine into the smoking cessation drug varenicline, the development of the blockbuster cardiovascular drug atorvastatin from the polyketide lovastatin, and the discovery of the proteasome inhibitor carfilzomib inspired by the natural polypeptide epoxomicin (**Fig. 1**). Such approaches, however, are greatly facilitated by an understanding of the binding mode to its protein target as well as the identification of key pharmacophores within the parent natural product (**Fig. 1**).

**Figure 1.**
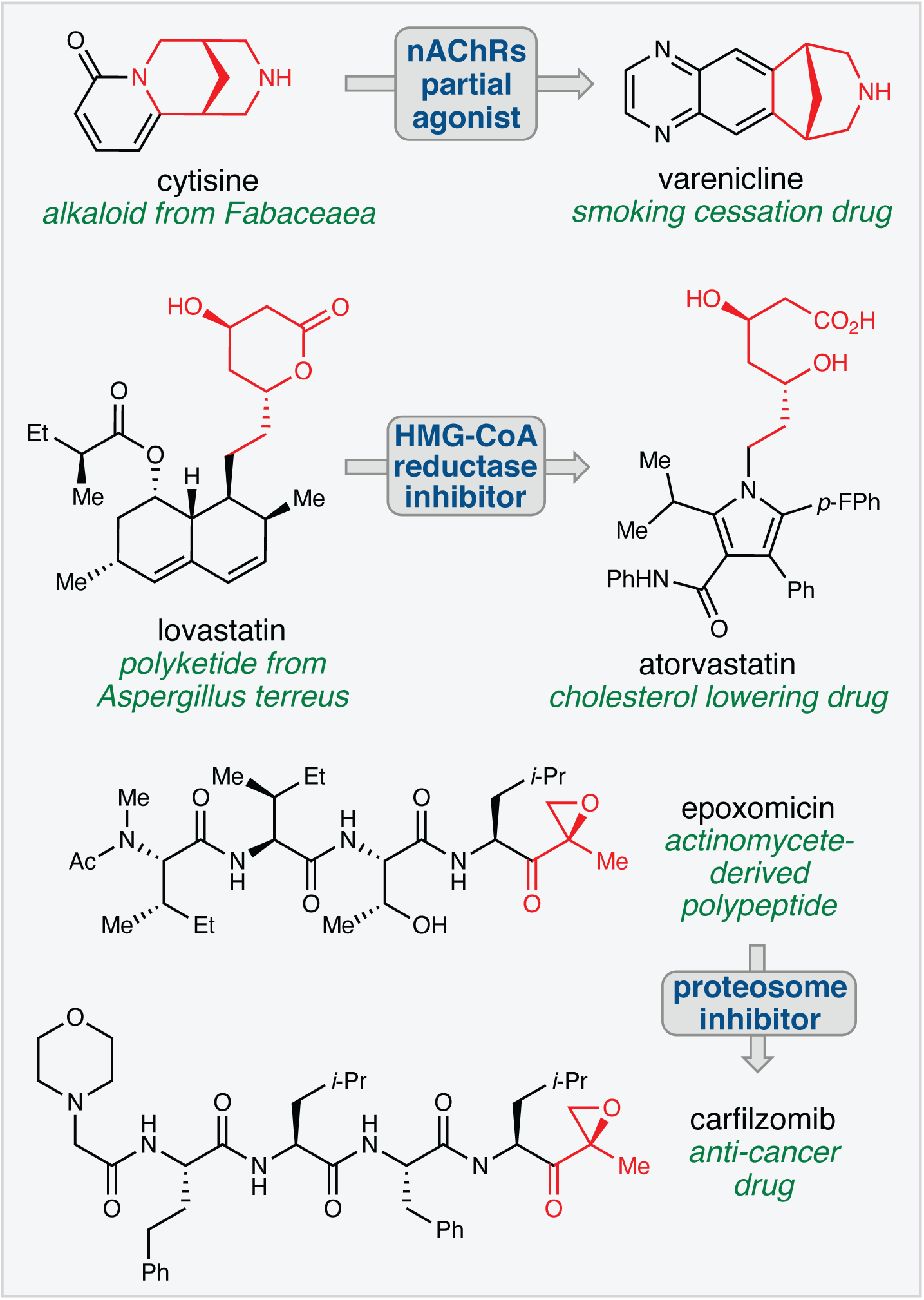
Translation of natural product function. Many successful drugs have recapitulated the function of natural products with fully-synthetic chemical scaffolds. Key pharmacophores are shown in red.

We recently discovered that the anti-cancer natural product nimbolide, a limonoid-type triterpene isolated from *Azadirachta indica* (neem), covalently reacts with an N-terminal cysteine (C8) within an intrinsically disordered region of the E3 ubiquitin ligase RNF114 in human breast cancer cells (Spradlin et al., 2019). Covalent targeting of RNF114 by nimbolide led to impaired ubiquitination of its endogenous substrate, tumor suppressor p21, through a nimbolide-dependent competition of the RNF114-substrate binding interaction, thus providing a potential mechanism for the anti-cancer effects of this natural product. The realization that nimbolide targeted a substrate recognition domain within RNF114 suggested that nimbolide could potentially be used as a novel recruiter of RNF114 for targeted protein degradation (TPD) applications. Consistent with this premise, we showed that a PROTAC formed by linking nimbolide to a Bromodomain and extraterminal domain (BET) inhibitor JQ1 led to proteasome-and RNF114-dependent degradation of BRD4 in cells. While TPD has arisen as a powerful drug discovery paradigm for tackling the undruggable proteome by targeting intracellular proteins for proteasomal degradation rather than classic inhibition, the lack of a broad range of E3 ligase recruiters represents a known limitation in this arena (Bondeson and Crews, 2017; Chamberlain and Hamann, 2019; Lai and Crews, 2017). Indeed, while over 600 different E3 ligases have been annotated, only a small handful of these potential targets have succumbed to the PROTAC strategy. Discovering additional, more synthetically tractable E3 ligase recruiters is thus an important topic in expanding the scope of TPD and may help to address resistance mechanisms (Bond et al., 2020; Ottis et al., 2019; Zeng et al., 2020), promote differing selectivity or kinetic profiles of degradation (Bondeson et al., 2018; Huang et al., 2018; Tong et al., 2020), and lead to cell-type or location-specific degradation.

Herein we realize the successful translation of the binding mode of nimbolide into a simple, easily manipulated covalent small molecule. Because the N-terminal region of RNF114 that includes C8 is intrinsically disordered, structure-guided ligand discovery and optimization was not possible thus requiring unbiased approaches to ligand discovery. Activity-based protein profiling (ABPP)-enabled covalent ligand screening has been previously used to discover novel covalent recruiters against E3 ligases RNF4 and DCAF16 (Ward et al., 2019; Zhang et al., 2019) and has also facilitated ligand discovery against cysteines targeted by covalently-acting natural products (Grossman et al., 2017). Using this technique, we were able to discover fragments that could be used to functionally replace nimbolide as the covalent E3 ligase recruitment module in fully functional degraders against several oncology targets.

## Results

### Covalent ligand screening against RNF114

To discover a fully synthetic covalent ligand that could access the same cysteine (C8) targeted by nimbolide on RNF114, we screened 318 cysteine-reactive chloroacetamide and acrylamide ligands in a gel-based competitive activity-based protein profiling (ABPP) assay, in which we competed ligands against binding of a rhodamine-functionalized cysteine-reactive iodoacetamide probe (IA-rhodamine) to pure RNF114 protein **(Fig. 2a, 2b, Supplementary Fig. 1 Supplementary Table 1)** (Bachovchin et al., 2010; Grossman et al., 2017; Ward et al., 2019). Through this screen, chloroacetamide EN219 emerged as the top hit, showing the greatest inhibition of IA-rhodamine binding to RNF114 **(Fig. 2c)**. Dose-response studies showed that EN219 interacted with RNF114 with a 50 % inhibitory concentration (IC50) of 470 nM **(Fig. 2d, 2e)**. EN219 also inhibited RNF114-mediated autoubiquitination and p21 ubiquitination *in vitro*, similarly to our previously observed findings with nimbolide **(Fig. 2f)**. Mapping the sites of EN219 covalent modification on pure RNF114 protein by LC-MS/MS also confirmed C8 as the primary site of modification **(Supplementary Fig. 2a)**.

**Figure 2.**
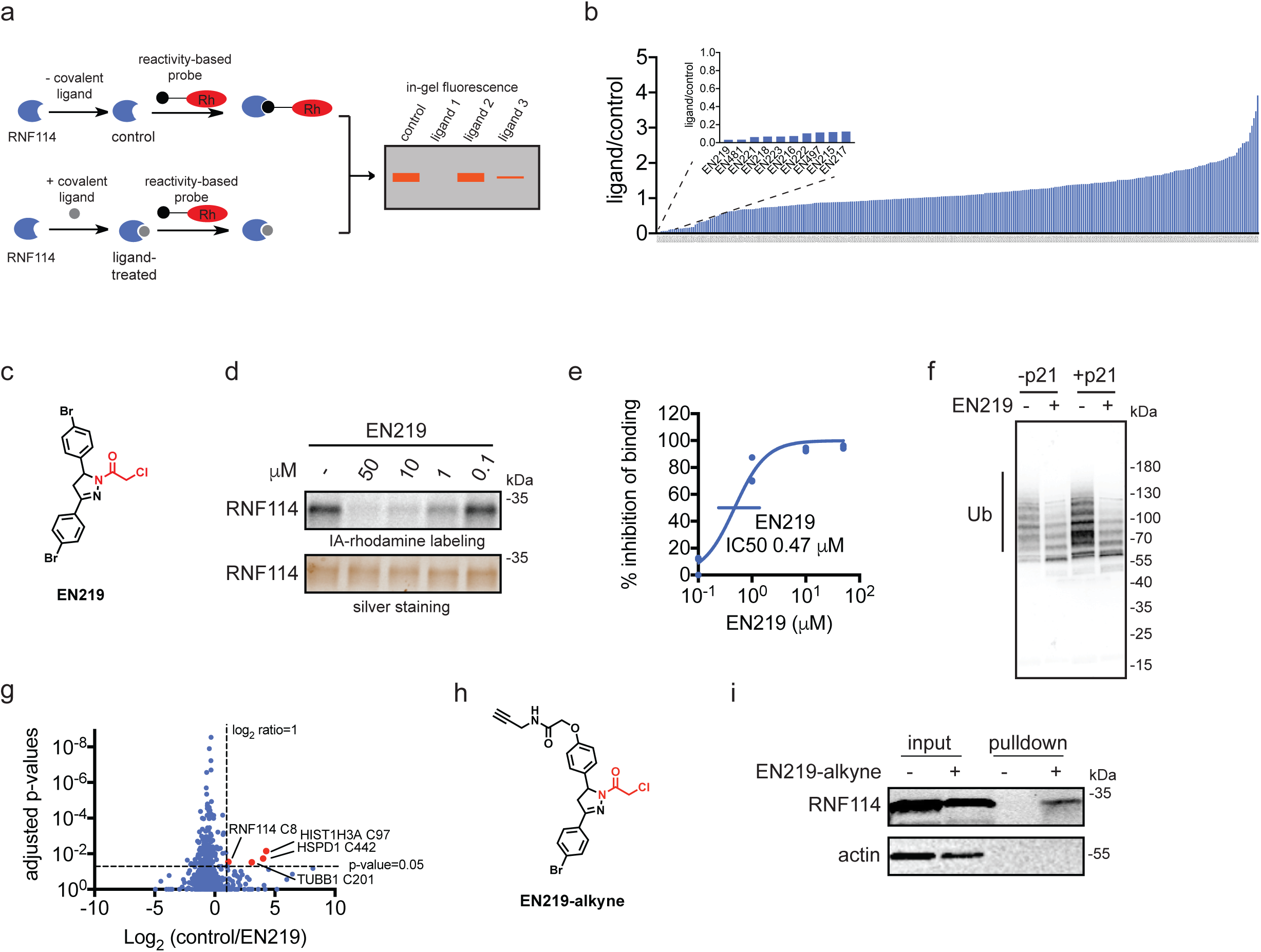
Covalent ligand screening against RNF114. **(a)** Gel-based ABPP assay for screening covalent ligands against IA-rhodamine probe binding to pure RNF114 protein. Loss of fluorescence indicates covalent ligand binding to a cysteine on RNF114. **(b)** Quantified results from gel-based ABPP screen of 318 cysteine-reactive acrylamides and chloroacetamides against IA-rhodamine labeling of RNF114. DMSO vehicle or covalent ligands (50 μM) were pre-incubated with pure RNF114 protein (0.1 μg) for 30 min prior to addition of IA-rhodamine (100 nM) for 30 min at room temperature. Proteins were separated by SDS/PAGE and in-gel fluorescence was quantified. Raw gel-based ABPP data shown in **Supplementary Fig. 1**. Structures of compounds screened can be found in **Supplementary Table 1**. Data expressed as ligand/control ratio of in-gel fluorescent intensity. Shown in the inlay are the ligand/control ratios for the top 10 hits. **(c)** Structure of top hit EN219 with chloroacetamide cysteine-reactive warhead in red. **(d)** Dose-response of EN219 interaction with RNF114 by competitive gel-based ABPP. DMSO vehicle or covalent ligands were pre-incubated with pure RNF114 protein (0.1 μg) for 30 min prior to addition of IA-rhodamine (100 nM) for 30 min at room temperature. Proteins were separated by SDS/PAGE and in-gel fluorescence was quantified. Gels were also silver-stained as a loading control. **(e)** Percent inhibition of IA-rhodamine binding to RNF114 in **(d)** was quantified and 50 % inhibitory concentration (IC50) was determined to be 0.47 μM. **(f)** RNF114-mediated autoubiquitination and p21 ubiquitination *in vitro*. DMSO vehicle or EN219 (50 μM) was incubated with pure RNF114 for 30 min prior to addition of PBS or p21, ATP, and FLAG-ubiquitin (Ub) for 60 min. Proteins were separated by SDS/PAGE and blotted for FLAG. Ubiquitinated-RNF114/p21 is noted. **(g)** IsoTOP-ABPP analysis of EN219 in 231MFP breast cancer cells. 231MFP cells were treated *in situ* with DMSO vehicle or EN219 (1 μM) for 90 min. Control and treated cell lysates were labeled with IA-alkyne (100 μM) for 1 h, after which isotopically light (control) or heavy (EN219-treated) biotin-azide bearing a TEV tag was appended by CuAAC. Proteomes were mixed in a 1:1 ratio, probe-labeled proteins were enriched with avidin and digested with trypsin, and probe-modified peptides were eluted by TEV protease and analyzed by LC-MS/MS. Average light-to-heavy (control/EN219) ratios and adjusted p-values were quantified for probe-modified peptides that were present in two out of three biological replicates and plotted. Shown in red are protein and modified cysteine site for peptides that showed >2-fold control/EN219 ratio with adjusted p-value <0.05. Full data can be found in **Supplementary Table 2. (h)** Structure of alkyne-functionalized EN219 probe. **(i)** EN219-alkyne pulldown of RNF114. 231MFP cells were treated with DMSO vehicle or EN219-alkyne (50 μM) for 90 min. Biotin-azide was appended to probe-labeled proteins by CuAAC and probe-labeled proteins were avidin enriched and blotted for RNF114 and negative control actin. An aliquot of input proteome from vehicle-and EN219-treated cell lysates was also subjected to blotting as an input control. Data shown in **(e)** are average values and individual replicate values. Data shown are from n=1 in **(b)** and n=3 for **(d-f, i)** biological replicate(s) per group. Gels shown in **(d, i)** are representative gels from n=3 biological replicates per group. This figure is related to **Supplementary Figures 1-2**.

To ascertain the selectivity of EN219, we next mapped the proteome-wide cysteine-reactivity of EN219 *in situ* in 231MFP breast cancer cells by competitive isotopic tandem orthogonal proteolysis-ABPP (isoTOP-ABPP) using previously well-validated methods (Backus et al., 2016; Grossman et al., 2017; Spradlin et al., 2019; Wang et al., 2014; Weerapana et al., 2010). Cells were treated *in situ* with vehicle or EN219 and cells were subsequently lysed and labeled with an alkyne-functionalized iodoacetamide probe (IA-alkyne), followed by attachment of isotopically light or heavy, Tobacco Etch Virus (TEV)-cleavable biotin-azide enrichment handles through copper-catalyzed azide-alkyne cycloaddition (CuAAC). Probe-labeled proteins were enriched by avidin, digested with trypsin, and probe-modified peptides were eluted by TEV protease and analyzed by liquid chromatography-mass spectrometry (LC-MS/MS). Only 4 proteins showed >2-fold light versus heavy or control versus EN219-treated probe-modified peptide ratios with adjusted p-value <0.05 out of 686 probe-modified cysteines quantified. These proteins were RNF114 C8, TUBB1 C201, HSPD1 C442, and HIST1H3A C97, among which RNF114 was the only E3 ligase **(Fig. 2g, Supplementary Table 2)**.

To further investigate the selectivity of EN219 and to confirm EN219 engagement of RNF114 in cells, we also synthesized an alkyne-functionalized EN219 probe (EN219-alkyne) **(Fig. 2h)**. We performed Tandem mass-tagging (TMT)-based quantitative proteomic profiling to identify proteins that were enriched by EN219-alkyne *in situ* labeling of 231MFP cells and were competed by EN219 *in situ* treatment **(Supplementary Fig. 2b Supplementary Table 3)**. While we did not identify RNF114 in this proteomics experiment, likely due to its relatively low abundance, we did identify 7 additional potential off-targets of EN219 that showed >3-fold competition with adjusted p-value <0.05 against EN219-alkyne labeling in cells **(Supplementary Fig. 2b)**. Again, none of these 7 potential EN219 off-targets were E3 ligases. Collectively, our results suggested that EN219 was a moderately selective covalent ligand against C8 of RNF114. While we did not observe RNF114 by TMT-based proteomic experiments, RNF114 was clearly enriched from 231MFP cells treated with EN219-alkyne *in situ* compared to vehicle-treated controls after CuAAC-mediated appendage of biotin-azide and subsequent avidin-pulldown and RNF114 blotting **(Fig. 2i)**.

### EN219-Based RNF114 Recruiter in TPD Applications

We next tested whether this fully synthetic RNF114 covalent ligand, EN219, could be exploited for TPD applications. We had previously demonstrated that nimbolide could be linked to the BET inhibitor ligand JQ1 to selectively degrade BRD4. Thus, we benchmarked our EN219 RNF114 recruiter by linking EN219 to JQ1 with three different linkers—ML 2-14, ML 2-31, and ML 2-32 with C4 alkyl, C7 alkyl, and polyethylene glycol (PEG) 4 linkers, respectively **(Fig. 3a; Supplementary Fig. 3a)**. Similar to nimbolide-based degraders, ML 2-14 with the shortest linker showed the most robust degradation of BRD4 in 231MFP breast cancer cells compared to ML 2-31 and ML 2-32 with longer linkers, with 50 % degradation concentration (DC50) values of 36 and 14 nM for the long and short isoforms of BRD4, respectively **(Fig. 3b, 3c; Supplementary Fig. 3b 3c)** (Spradlin et al., 2019). EN219 treatment alone did not affect BRD4 levels, compared to ML 2-14 **(Supplementary Fig. 4a)**. This ML 2-14 mediated degradation was fully averted by pre-treating cells with the proteasome inhibitor bortezomib as well as the E1 activating enzyme inhibitor TAK-243 **(Fig. 3d-3e, Supplementary Fig. 4b, 4c)**.

**Figure 3.**
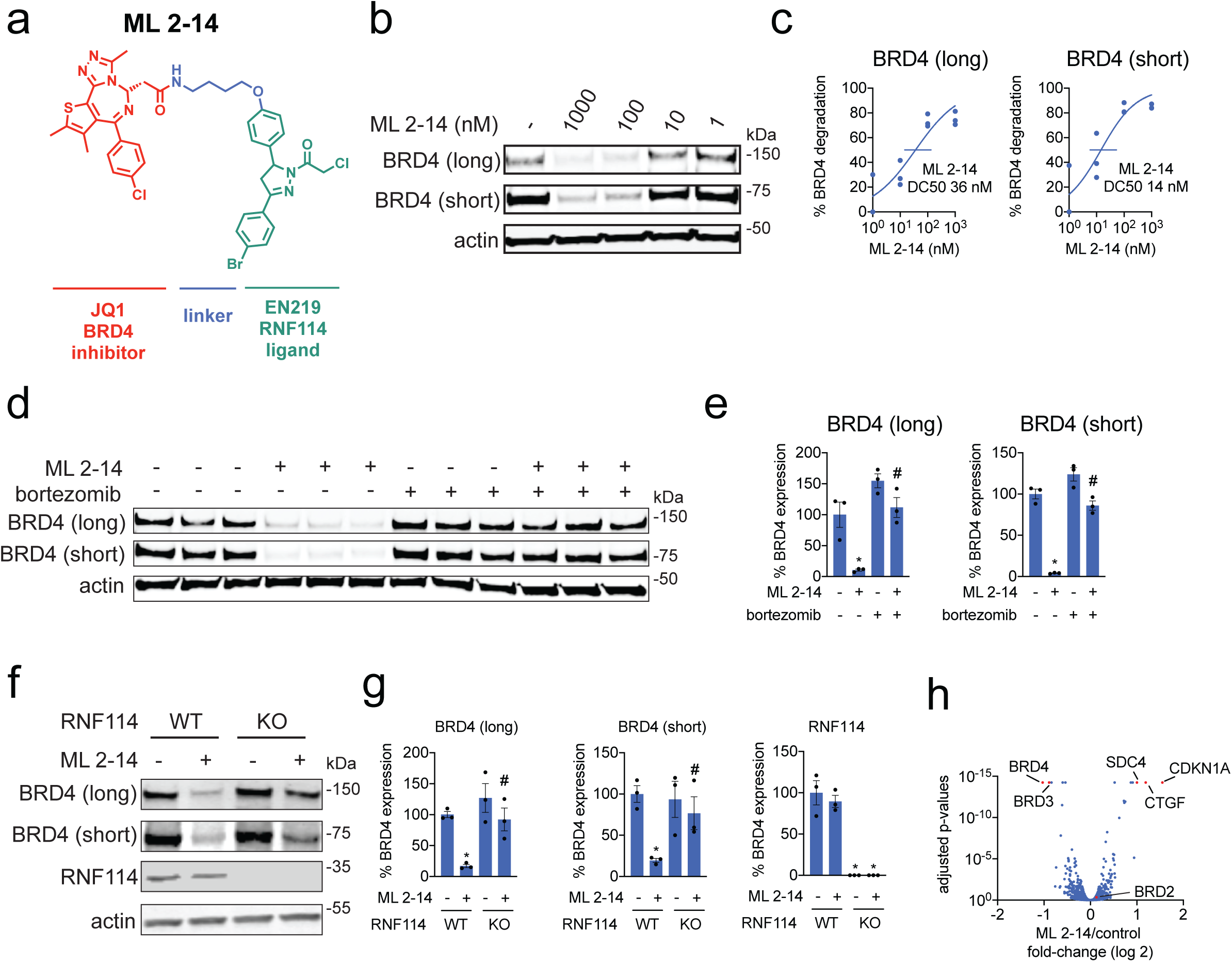
EN219-based BRD4 degrader. **(a)** Structure of ML 2-14, an EN219-based BRD4 degrader linking EN219 to BET inhibitor JQ1. **(b)** Degradation of BRD4 by ML 2-14. 231MFP cells were treated with DMSO vehicle or ML 2-14 for 8 h and the long and short isoforms of BRD4 and loading control actin were detected by Western blotting. **(c)** Percentage of BRD4 degradation quantified from **(b)** show 50 % degradation concentrations of 36 and 14 nM for the long and short isoforms of BRD4, respectively. **(d)** Proteasome-dependent degradation of BRD4 by ML 2-14. 231MFP cells were treated with DMSO vehicle or with proteasome inhibitor bortezomib (1 μM) 30 min prior to DMSO vehicle or ML 2-14 (100 nM) treatment for 8 h. BRD4 and loading control actin levels were detected by Western blotting. **(e)** Quantification of BRD4 levels from **(d). (f)** Degradation of BRD4 in RNF114 wild-type (WT) or knockout (KO) HAP1 cells. RNF114 WT or KO HAP1 cells were treated with DMSO vehicle or ML 2-14 (1 μM) for 16 h and BRD4, RNF114, and loading control actin levels were detected by Western blotting. **(g)** Quantification of data from **(f). (h)** TMT-based quantitative proteomic data showing ML 2-14-mediated protein level changes in 231MFP cells. 231MFP cells were treated with DMSO vehicle or ML 2-14 for 8 h. Shown are average TMT ratios for all tryptic peptides identified with at least 2 unique peptides. Shown in red are proteins that showed >2-fold increased or decreased levels with ML 2-14 treatment with adjusted p-values <0.05. Data in **(h)** are from n=3 biological replicates/group. Data shown in **(c)** are average and individual replicate values. Data shown in **(e, g)** are averages ± sem values and also individual biological replicate values. Blots shown in **(b, d, f)** are representative of n=3 biological replicates/group. This figure is related to **Supplementary Figures 3-4**.

**Figure 4.**
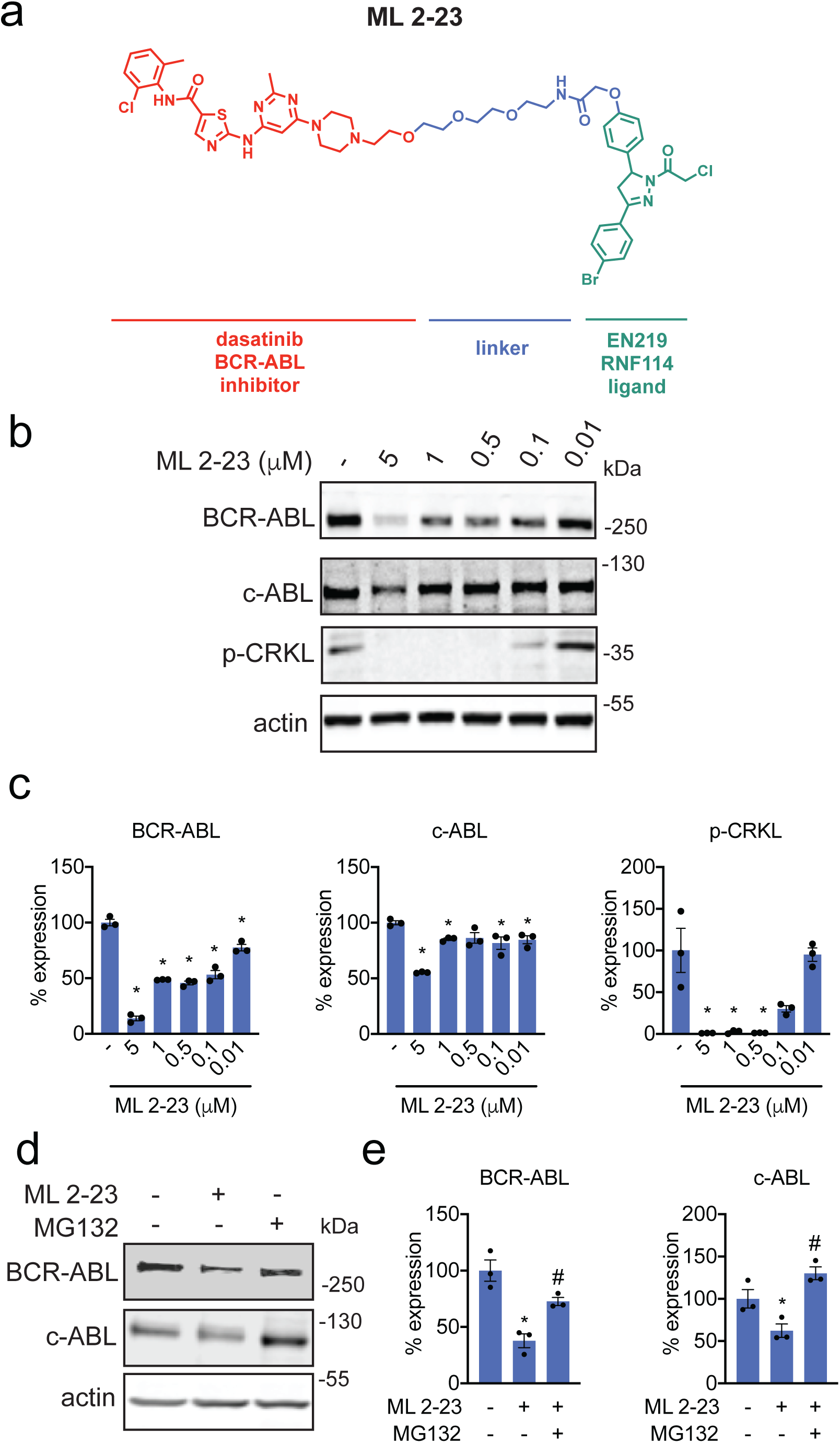
EN219-based ABL degrader. **(a)** Structure of ML 2-23, an EN219-based ABL degrader linking EN219 to ABL inhibitor dasatinib. **(b)** Degradation of BCR-ABL and c-ABL by ML 2-23. K562 cells were treated with DMSO vehicle or ML 2-23 for 16 h and BCR-ABL, c-ABL, p-CRKL, and loading control actin were detected by Western blotting. **(c)** Percentage of BCR-ABL, c-ABL, and p-CRKL expression quantified from **(b). (d)** Proteasome-dependent degradation of BCR-ABL and c-ABL by ML 2-23. K562 cells were treated with DMSO vehicle or with proteasome inhibitor MG132 (5 μM) 30 min prior to DMSO vehicle or ML 2-23 (1 μM) treatment for 12 h. BCR-ABL, c-ABL, and loading control actin levels were detected by Western blotting. **(e)** Quantification of BCR-ABL and c-ABL levels from **(d)**. Data shown in **(c, e)** are averages ± sem values and also individual biological replicate values. Blots shown in **(b, d)** are representative of n=3 biological replicates/group. This figure is related to **Supplementary Figure 4**.

To further validate that ML 2-14 degradation of BRD4 was driven through RNF114, we showed that BRD4 degradation in HAP1 cells was significantly attenuated in RNF114 knockout cells compared to wild-type counterparts **(Fig. 3f-3g)**. TMT-based quantitative proteomic profiling of ML 2-14-mediated protein expression changes showed selective degradation of BRD3 and BRD4, but not BRD2. We also observed stabilization of two known or putative RNF114 substrates, including the tumor suppressor CDKN1A (p21) and CTGF **(Fig. 3h, Supplementary Table 4)** (Han et al., 2013; Spradlin et al., 2019).

To further demonstrate the utility of our newly discovered fully synthetic RNF114 recruiter EN219 in degrading other more challenging protein targets, we synthesized a degrader linking EN219 to the BCR-ABL inhibitor dasatinib, ML 2-23 and ML 2-22, bearing a longer PEG3 linker and a shorter C3 alkyl linker, respectively **(Fig. 4a, Supplementary Fig. 5a)**. For this particular target, ML 2-23 with the longer linker showed more robust degradation of BCR-ABL in K562 leukemia cells compared to ML 2-22, consistent with previously observed structure-activity relationships of nimbolide-based BCR-ABL degraders **(Fig. 4b-4c, Supplementary Fig. 5b)** (Tong et al., 2020). Consistent with ML 2-23 engaging BCR-ABL in cells, we observed inhibition of CRKL phosphorylation, a downstream substrate of BCR-ABL signaling **(Fig. 4b-4c)**. Interestingly, EN219 showed preferential degradation of BCR-ABL compared to c-ABL, compared to several previous BCR-ABL degraders utilizing cereblon or VHL recruiters that showed opposite selectivity **(Fig. 4b-4c)** (Burslem et al., 2019; Lai et al., 2016). This preferential degradation was also observed with the equivalent nimbolide-based degrader ^9^. While rescue experiments with proteasome inhibitors proved challenging due to the cytotoxicity of proteasome inhibitors at the long timepoints required for robust BCR-ABL degradation, we observed significant rescue of early stage ML 2-23-mediated BCR-ABL and c-ABL degradation with pre-treatment of K562 cells with the proteasome inhibitor MG132 at a shorter time point **(Fig. 4d, 4e)**.

## Discussion

We previously discovered that the natural product nimbolide targets a predicted intrinsically disordered cysteine in RNF114 at a substrate recognition site (Spradlin et al., 2019). Here we report EN219 as a moderately selective covalent ligand that exploits this same binding modality. We show that EN219 can be linked to the BET inhibitor JQ1 to degrade BRD4 in a nimbolide-sensitive and proteasome- and RNF114-dependent manner. We further show that EN219 can be linked to the kinase inhibitor dasatinib to selectively degrade BCR-ABL over c-ABL in a proteasome-dependent manner in leukemia cells. Thus, EN219 represents a fully synthetic and more tunable chemical scaffold for targeting RNF114 for TPD applications relative to the complex natural product from which it was inspired. These findings also further highlight that moderately selective cysteine-targeting ligands can still lead to robust protein degraders (Ward et al., 2019; Zhang et al., 2019). While EN219 represents a promising initial chemical scaffold for RNF114 recruitment, future medicinal chemistry efforts will be required to further optimize the potency and metabolic stability of RNF114 recruiters, but its synthetic tractability compared to nimbolide makes this a realistic endeavor.

Overall, our results highlight the utility of chemoproteomic platforms for discovering new chemical scaffolds that can be used as E3 ligase recruiters for TPD applications. Moreover, these studies further speak to the power of unbiased chemoproteomic approaches in mimicking the function of covalently acting natural products, particularly those in which structural binding information is unavailable (Nomura and Maimone, 2019).

## Methods

### Chemicals

Covalent ligands screened against RNF114 were purchased from Enamine LLC, including EN219. Structures of compounds screened can be found in **Supplementary Table 1**. See Supplementary Information for synthetic methods and characterization for EN219 and degraders.

### Cell culture

The 231MFP cells were obtained from Prof. Benjamin Cravatt and were generated from explanted tumor xenografts of MDA-MB-231 cells as previously described (Jessani et al., 2004). The 231MFP cells were cultured in L15 medium containing 10% FBS and maintained at 37 °C with 0% CO_2_. K562 chronic myeloid leukemia cell lines were purchased from ATCC. The K562 cells were cultured in Iscove’s Modified Dμlbecco’s Medium containing 10% FBS and maintained at 37 °C with 5% CO_2_. HAP1 RNF114 wild-type and knockout cell lines were purchased from Horizon Discovery. The RNF114 knockout cell line was generated by CRISPR/Cas9 to contain a frameshift mutation in a coding exon of RNF114. HAP1 cells were grown in Iscove’s Modifed Dμlbecco’s Medium in the presence of 10% FBS and penicillin/streptomycin.

### Cell-based degrader assays

For assaying degrader activity, cells were seeded (500,000 cells for 231MFP and HAP1 cells, 1,000,000 for K562 cells) into a 6 cm tissue culture dish (Corning) in 2.0−2.5 mL of media and allowed to adhere overnight for 231MFP and HAP1 cells. The following morning, media was replaced with complete media containing the desired concentration of compound diluted from a 1,000 x stock in DMSO. For rescue studies, the cells were pre-treated with proteasome inhibitors or E1 activating enzyme inhibitors 30 min prior to the addition of DMSO or degrader compounds. At the specified time point, cells were washed once with PBS on ice, before addition of 120 μL of lysis buffer (20 mM Tris-HCl at pH 7.5, 150 mM NaCl, 1 mM Na_2_EDTA, 1 mM EGTA, 1% Triton, 2.5 mM sodium pyrophosphate, 1 mM beta-glycerophosphate, 1 mM Na_3_VO_4_, 1 µg/ml leupeptin) with Complete Protease Inhibitor Cocktail (Sigma) was added. The cells were incubated in lysis buffer for 5 min before scraping and transferring to microcentrifuge tubes. The lysates were then frozen at −80 °C or immediately processed for Western blotting. To prepare for Western blotting, the lysates were cleared with a 20,000g spin for 10 min and the resulting supernatant protein concentrations were quantified via BCA assay. The lysates were normalized by dilution with PBS to match the lowest concentration lysate, and the appropriate amount of 4 x Laemmli’s reducing buffer was added.

### Gel-Based ABPP

Gel-Based ABPP methods were performed as previously described (Ward et al., 2019). Pure recombinant human RNF114 was purchased from Boston Biochem (K-220). RNF114 (0.25 μg) was diluted into 50 μL of PBS and 1 μL of either DMSO (vehicle) or covalently acting small molecule to achieve the desired concentration. After 30 min at room temperature, the samples were treated with 250 nM of tetramethylrhodamine-5-iodoacetamide dihydroiodide (IA-Rhodamine) (Setareh Biotech, 6222, prepared in anhydrous DMSO) for 1 h at room temperature. Incubations were quenched by diluting the incubation with 20 μL of 4 x reducing Laemmli SDS sample loading buffer (Alfa Aesar) and heated at 90 °C for 5 min. The samples were separated on precast 4−20% Criterion TGX gels (Bio-Rad Laboratories, Inc.). Fluorescent imaging was performed on a ChemiDoc MP (Bio-Rad Laboratories, Inc.). Inhibition of target labeling was assessed by densitometry using ImageJ.

### EN219-alkyne probe labeling *in situ* and pulldown studies

Experiments were performed following an adaption of a previously described protocol (Thomas et al., 2017). The 231MFP cells were treated with either DMSO vehicle or 50 μM EN219-alkyne probe for 90 min. Cells were collected in PBS and lysed by sonication. For preparation of Western blotting samples, the lysate (1 mg of protein in 500 μl) was aliquoted per sample and then the following were added: 10 μl of 5 mM biotin picolylazide (900912 Sigma-Aldrich) and 50 μl of click reaction mix (three parts TBTA 5 mM TBTA in butanol:DMSO (4:1, v/v), one part 50 mM Cu(II)SO_4_ solution and one part 50 mM TCEP). Samples were incubated for 1 h at room temperature with gentle agitation. After CuAAC, proteomes were precipitated by centrifugation at 6,500 *g* and washed twice in ice-cold methanol (500 μl). The samples were spun in a prechilled (4 °C) centrifuge at 6,500 *g* for 4 min allowing for aspiration of excess methanol and subsequent reconstitution of protein pellet in 250 μl PBS containing 1.2% SDS by probe sonication. Then the proteome was denatured at 90 °C for 5 min, the insoluble components were precipitated by centrifugation at 6,500*g* and soluble proteome was diluted in 1.2 ml PBS (the final concentration of SDS in the sample was 0.2%) to a total volume of 1450 μl, with 50 μl reserved as input. Then 85 μl of prewashed 50% streptavidin agarose bead slurry was added to each sample and samples were incubated overnight at room temperature with gentle agitation. Supernatant was aspirated from each sample after spinning beads at 6,500 *g* for 2 min at room temperature. Beads were transferred to spin columns and washed three times with PBS. To elute, beads were boiled 5 min in 50 μl LDS sample buffer. Eluents were collected after centrifugation and analyzed by immunoblotting. The resulting samples were also analyzed as described below for TMT-based quantitative proteomic profiling.

### LC–MS/MS analysis of pure RNF114 EN219 modification

Purified RNF114 (10 μg) in 50μl PBS was incubated 30 min at room temperature either with DMSO vehicle or EN219 (50 μM). The DMSO control was then treated with light iodoacetamide while the compound treated sample was incubated with heavy iodoacetamide for 1h each at room temperature (200 μM final concentration, Sigma-Aldrich, 721328). The samples were precipitated by addition of 12.5 µl of 100% (w/v) trichloroacetic acid and the treated and control groups were combined pairwise, before cooling to −80 °C for 1h. The combined sample was then spun for at max speed for 10 min at 4 °C, supernatant was carefully removed and the sample was washed with ice-cold 0.01 M HCl/90% acetone solution. The pellet was resuspended in 4M urea containing 0.1% Protease Max (Promega Corp. V2071) and diluted in 40 mM ammonium bicarbonate buffer. The samples were reduced with 10 mM TCEP at 60 °C for 30min. The sample was then diluted 50% with PBS before sequencing grade trypsin (1 μg per sample, Promega Corp, V5111) was added for an overnight incubation at 37 °C. The next day, the sample was centrifuged at 13,200 rpm for 30 min. The supernatant was transferred to a new tube and acidified to a final concentration of 5% formic acid and stored at −80 °C until mass spectrometry analysis.

### IsoTOP-ABPP

IsoTOP-ABPP studies were done as previously reported (Spradlin et al., 2019). Cells were lysed by probe sonication in PBS and protein concentrations were measured by BCA assay35. For in situ experiments, cells were treated for 90 min with either DMSO vehicle or covalently acting small molecμle (from 1,000× DMSO stock) before cell collection and lysis. Proteomes were subsequently labeled with *N*-5-Hexyn-1-yl-2-iodoacetamide (IA-alkyne) labeling (100 μM) for 1 h at room temperature. CuAAC was used by sequential addition of tris(2-carboxyethyl) phosphine (1 mM, Sigma), tris[(1-benzyl-1H-1,2,3-triazol-4-yl)methyl]amine (34 μM, Sigma), copper(II) sμlfate (1 mM, Sigma) and biotin-linker-azide—the linker functionalized with a tobacco etch virus (TEV) protease recognition sequence as well as an isotopically light or heavy valine for treatment of control or treated proteome, respectively. After CuAAC, proteomes were precipitated by centrifugation at 6,500*g*, washed in ice-cold methanol, combined in a 1:1 control:treated ratio, washed again, then denatured and resolubilized by heating in 1.2% SDS–PBS to 80 °C for 5 min. Insoluble components were precipitated by centrifugation at 6,500*g* and soluble proteome was diluted in 5 ml 0.2% SDS–PBS. Labeled proteins were bound to avidin-agarose beads (170 μl resuspended beads per sample, Thermo Pierce) while rotating overnight at 4 °C. Bead-linked proteins were enriched by washing three times each in PBS and water, then resuspended in 6 M urea and PBS (Sigma), and reduced in TCEP (1 mM, Sigma), alkylated with iodoacetamide (18 mM, Sigma), before being washed and resuspended in 2 M urea and trypsinized overnight with 0.5 μg μl^−1^ sequencing grade trypsin (Promega). Tryptic peptides were eluted off. Beads were washed three times each in PBS and water, washed in TEV buffer solution (water, TEV buffer, 100 μM dithiothreitol) and resuspended in buffer with Ac-TEV protease and incubated overnight. Peptides were diluted in water and acidified with formic acid (1.2 M, Spectrum) and prepared for analysis.

### Mass spectrometry analysis

Total peptides from TEV protease digestion for isoTOP-ABPP or tryptic peptides for mapping EN219 site of modification on pure RNF114 were pressure loaded onto 250 mm tubing packed with Aqua C18 reverse phase resin (Phenomenex no. 04A-4299), which was previously equilibrated on an Agilent 600 series high-performance liquid chromatograph using the gradient from 100% buffer A to 100% buffer B over 10 min, followed by a 5 min wash with 100% buffer B and a 5 min wash with 100% buffer A. The samples were then attached using a MicroTee PEEK 360 μm fitting (Thermo Fisher Scientific no. p-888) to a 13 cm laser pμlled column packed with 10 cm Aqua C18 reverse-phase resin and 3 cm of strong-cation exchange resin for isoTOP-ABPP studies. Samples were analyzed using an Q Exactive Plus mass spectrometer (Thermo Fisher Scientific) using a five-step Mμltidimensional Protein Identification Technology (MudPIT) program, using 0, 25, 50, 80 and 100% salt bumps of 500 mM aqueous ammonium acetate and using a gradient of 5–55% buffer B in buffer A (buffer A: 95:5 water:acetonitrile, 0.1% formic acid; buffer B 80:20 acetonitrile:water, 0.1% formic acid). Data were collected in datadependent acquisition mode with dynamic exclusion enabled (60 s). One fμll mass spectrometry (MS^1^) scan (400–1,800 mass-to-charge ratio (m/z)) was followed by 15 MS^2^ scans of the nth most abundant ions. Heated capillary temperature was set to 200 °C and the nanospray voltage was set to 2.75 kV, as previously described.

Data were extracted in the form of MS^1^ and MS^2^ files using Raw Extractor v.1.9.9.2 (Scripps Research Institute) and searched against the Uniprot human database using ProLuCID search methodology in IP2 v.3 (Integrated Proteomics Applications, Inc.) (Xu et al., 2015). Probe-modified cysteine residues were searched with a static modification for carboxyaminomethylation (+57.02146) and up to two differential modifications for methionine oxidation and either the light or heavy TEV tags (+464.28596 or +470.29977, respectively) or the mass of the EN219 adduct. Peptides were required to be fully tryptic peptides and to contain the TEV modification. ProLUCID data was filtered through DTASelect to achieve a peptide false-positive rate below 5%. Only those probe-modified peptides that were evident across two out of three biological replicates were interpreted for their isotopic light to heavy ratios. Light versus heavy isotopic probe-modified peptide ratios are calculated by taking the mean of the ratios of each replicate paired light vs. heavy precursor abundance for all peptide spectral matches (PSM) associated with a peptide. The paired abundances were also used to calculate a paired sample t-test p-value in an effort to estimate constancy within paired abundances and significance in change between treatment and control. P-values were corrected using the Benjamini/Hochberg method.

### TMT-based quantitative proteomic profiling

TMT proteomic profiling was performed as previously described (Spradlin et al., 2019).

### RNF114 ubiquitination assay

Recombinant Myc-Flag-RNF114 proteins were purchased from Origene (Origene Technologies Inc., TP309752) or were purified as described previously(Spradlin et al., 2019). For in vitro auto-ubiquitination assay, 0.2 μg of RNF114 in 25 μl of TBS was pre-incubated with DMSO vehicle or the covalently acting compound for 30 min at room temperature. Subsequently, 0.1 μg of UBE1 (Boston Biochem. Inc., E-305), 0.1 μg UBE2D1 (Boston Bichem. Inc., e2-615), 5 μg of Flag-ubiquitin (Boston Bichem. Inc., u-120) in a total volume of 25 μl Tris buffer containing 2 mM ATP, 10 mM DTT and 10 mM MgCl_2_ were added to achieve a final volume of 50 μl. For substrate-protein ubiquitination assays, 0.1 μg of p21 (Origene) was added at this stage. The mixture was incubated at 37 °C with agitation for 1.5 h. Then, 20 μl of Laemmli’s buffer was added to quench the reaction and proteins were analyzed by Western blot assay.

### Western blotting

Antibodies to RNF114 (Millipore Sigma, HPA021184), c-ABL (Santa Crus, 24-11), p-CRKL (Tyr207, Cell Signaling Technology, 3181), GAPDH (Proteintech Group Inc., 60004-1-Ig), BRD4 (Abcam plc, Ab128874), and beta-actin (Proteintech Group Inc., 6609-1-Ig) were obtained from various commercial sources and dilutions were prepared per the recommended manufacturers’ procedures. Proteins were resolved by SDS– PAGE and transferred to nitrocellulose membranes using the iBlot system (Invitrogen). Blots were blocked with 5% BSA in Tris-buffered saline containing Tween 20 (TBST) solution for 1 h at room temperature, washed in TBST and probed with primary antibody diluted in diluent, as recommended by the manufacturer, overnight at 4 °C. Following washes with TBST, the blots were incubated in the dark with secondary antibodies purchased from Ly-Cor and used at 1:10,000 dilution in 5% BSA in TBST at room temperature. Blots were visualized using an Odyssey Li-Cor scanner after additional washes. If additional primary antibody incubations were required, the membrane was stripped using ReBlot Plus Strong Antibody Stripping Solution (EMD Millipore, 2504), washed and blocked again before being re-incubated with primary antibody. Blots were quantified and normalized to loading controls using Image J.

## Data Availability Statement

The datasets generated during and/or analyzed during the current study are available from the corresponding author on reasonable request.

## Code Availability Statement

Data processing and statistical analysis algorithms from our lab can be found on our lab’s Github site: https://github.com/NomuraRG, and we can make any further code from this study available at reasonable request.

## Acknowledgement

We thank the members of the Nomura Research Group and Novartis Institutes for BioMedical Research for critical reading of the manuscript. This work was supported by Novartis Institutes for BioMedical Research and the Novartis-Berkeley Center for Proteomics and Chemistry Technologies (NB-CPACT) for all listed authors. This work was also supported by the Nomura Research Group and the Mark Foundation for Cancer Research and Chordoma Foundation ASPIRE Award for DKN, ML, JNS. This work was also supported by grants from the National Institutes of Health (R01CA240981 for DKN, ML, JNS and F31CA239327 for JNS).

## Author Contributions

ML, DKN conceived the project and wrote the paper. ML, JNS, DKN, JAT, MS, JMK, SMB provided intellectual contributions and insights into project direction. ML, JNS, SMB, MS, DKN designed the experiments. ML, JNS, SMB, DKN performed experiments and analyzed data. YI, JAT, JMK, LM, MS, SMB, MDJ, XL, WF, TJM, DKN edited the paper.

## References

Bachovchin, D.A., Ji, T., Li, W., Simon, G.M., Blankman, J.L., Adibekian, A., Hoover, H., Niessen, S., and Cravatt, B.F. (2010). Superfamily-wide portrait of serine hydrolase inhibition achieved by library-versus-library screening. Proc. Natl. Acad. Sci. U.S.A. 107, 20941–20946.

Backus, K.M., Correia, B.E., Lum, K.M., Forli, S., Horning, B.D., González-Páez, G.E., Chatterjee, S., Lanning, B.R., Teijaro, J.R., Olson, A.J., et al. (2016). Proteome-wide covalent ligand discovery in native biological systems. Nature 534, 570–574.

Bond, M.J., Chu, L., Nalawansha, D.A., Li, K., and Crews, C. (2020). Targeted Degradation of Oncogenic KRASG12C by VHL-recruiting PROTACs.

Bondeson, D.P., and Crews, C.M. (2017). Targeted Protein Degradation by Small Molecules. Annual Review of Pharmacology and Toxicology 57, 107–123.

Bondeson, D.P., Smith, B.E., Burslem, G.M., Buhimschi, A.D., Hines, J., Jaime-Figueroa, S., Wang, J., Hamman, B.D., Ishchenko, A., and Crews, C.M. (2018). Lessons in PROTAC Design from Selective Degradation with a Promiscuous Warhead. Cell Chemical Biology 25, 78-87.e5.

Burslem, G.M., Schultz, A.R., Bondeson, D.P., Eide, C.A., Savage Stevens, S.L., Druker, B.J., and Crews, C.M. (2019). Targeting BCR-ABL1 in Chronic Myeloid Leukemia by PROTAC-Mediated Targeted Protein Degradation. Cancer Res. 79, 4744–4753.

Chamberlain, P.P., and Hamann, L.G. (2019). Development of targeted protein degradation therapeutics. Nat. Chem. Biol. 15, 937–944.

Grossman, E.A., Ward, C.C., Spradlin, J.N., Bateman, L.A., Huffman, T.R., Miyamoto, D.K., Kleinman, J.I., and Nomura, D.K. (2017). Covalent Ligand Discovery against Druggable Hotspots Targeted by Anti-cancer Natural Products. Cell Chem Biol 24, 1368-1376.e4.

Han, J., Kim, Y.-L., Lee, K.-W., Her, N.-G., Ha, T.-K., Yoon, S., Jeong, S.-I., Lee, J.-H., Kang, M.-J., Lee, M.-G., et al. (2013). ZNF313 is a novel cell cycle activator with an E3 ligase activity inhibiting cellular senescence by destabilizing p21(WAF1.). Cell Death Differ. 20, 1055–1067.

Huang, H.-T., Dobrovolsky, D., Paulk, J., Yang, G., Weisberg, E.L., Doctor, Z.M., Buckley, D.L., Cho, J.-H., Ko, E., Jang, J., et al. (2018). A Chemoproteomic Approach to Query the Degradable Kinome Using a Multi-kinase Degrader. Cell Chemical Biology 25, 88-99.e6.

Jessani, N., Humphrey, M., McDonald, W.H., Niessen, S., Masuda, K., Gangadharan, B., Yates, J.R., Mueller, B.M., and Cravatt, B.F. (2004). Carcinoma and stromal enzyme activity profiles associated with breast tumor growth in vivo. Proc Natl Acad Sci U S A 101, 13756–13761.

Lai, A.C., and Crews, C.M. (2017). Induced protein degradation: an emerging drug discovery paradigm. Nat Rev Drug Discov 16, 101–114.

Lai, A.C., Toure, M., Hellerschmied, D., Salami, J., Jaime-Figueroa, S., Ko, E., Hines, J., and Crews, C.M. (2016). Modular PROTAC Design for the Degradation of Oncogenic BCR-ABL. Angew. Chem. Int. Ed. Engl. 55, 807–810.

Newman, D.J., and Cragg, G.M. (2016). Natural Products as Sources of New Drugs from 1981 to 2014. J. Nat. Prod. 79, 629–661.

Nomura, D.K., and Maimone, T.J. (2019). Target Identification of Bioactive Covalently Acting Natural Products. Curr. Top. Microbiol. Immunol. 420, 351–374.

Ottis, P., Palladino, C., Thienger, P., Britschgi, A., Heichinger, C., Berrera, M., Julien-Laferriere, A., Roudnicky, F., Kam-Thong, T., Bischoff, J.R., et al. (2019). Cellular Resistance Mechanisms to Targeted Protein Degradation Converge Toward Impairment of the Engaged Ubiquitin Transfer Pathway. ACS Chem. Biol. 14, 2215–2223.

Spradlin, J.N., Hu, X., Ward, C.C., Brittain, S.M., Jones, M.D., Ou, L., To, M., Proudfoot, A., Ornelas, E., Woldegiorgis, M., et al. (2019). Harnessing the anti-cancer natural product nimbolide for targeted protein degradation. Nat. Chem. Biol. 15, 747–755.

Thomas, J.R., Brittain, S.M., Lipps, J., Llamas, L., Jain, R.K., and Schirle, M. (2017). A Photoaffinity Labeling-Based Chemoproteomics Strategy for Unbiased Target Deconvolution of Small Molecule Drug Candidates. Methods Mol. Biol. 1647, 1–18.

Tong, B., Spradlin, J.N., Novaes, L.F.T., Zhang, E., Hu, X., Moeller, M., Brittain, S.M., McGregor, L.M., McKenna, J.M., Tallarico, J.A., et al. (2020). A Nimbolide-Based Kinase Degrader Preferentially Degrades Oncogenic BCR-ABL. BioRxiv 2020.04.02.022541.

Wang, C., Weerapana, E., Blewett, M.M., and Cravatt, B.F. (2014). A chemoproteomic platform to quantitatively map targets of lipid-derived electrophiles. Nat. Methods 11, 79–85.

Ward, C.C., Kleinman, J.I., Brittain, S.M., Lee, P.S., Chung, C.Y.S., Kim, K., Petri, Y., Thomas, J.R., Tallarico, J.A., McKenna, J.M., et al. (2019). Covalent Ligand Screening Uncovers a RNF4 E3 Ligase Recruiter for Targeted Protein Degradation Applications. ACS Chem. Biol. 14, 2430–2440.

Weerapana, E., Wang, C., Simon, G.M., Richter, F., Khare, S., Dillon, M.B.D., Bachovchin, D.A., Mowen, K., Baker, D., and Cravatt, B.F. (2010). Quantitative reactivity profiling predicts functional cysteines in proteomes. Nature 468, 790–795.

Xu, T., Park, S.K., Venable, J.D., Wohlschlegel, J.A., Diedrich, J.K., Cociorva, D., Lu, B., Liao, L., Hewel, J., Han, X., et al. (2015). ProLuCID: An improved SEQUEST-like algorithm with enhanced sensitivity and specificity. J Proteomics 129, 16–24.

Zeng, M., Xiong, Y., Safaee, N., Nowak, R.P., Donovan, K.A., Yuan, C.J., Nabet, B., Gero, T.W., Feru, F., Li, L., et al. (2020). Exploring Targeted Degradation Strategy for Oncogenic KRASG12C. Cell Chem Biol 27, 19-31.e6.

Zhang, X., Crowley, V.M., Wucherpfennig, T.G., Dix, M.M., and Cravatt, B.F. (2019). Electrophilic PROTACs that degrade nuclear proteins by engaging DCAF16. Nat. Chem. Biol. 15, 737–746.

