## Supplemental Methods and Figures for "Chemoproteomics-Enabled Ligand Screening Yields Covalent RNF114-Based Degraders that Mimic Natural Product Function"

### Synthetic Methods and Characterization

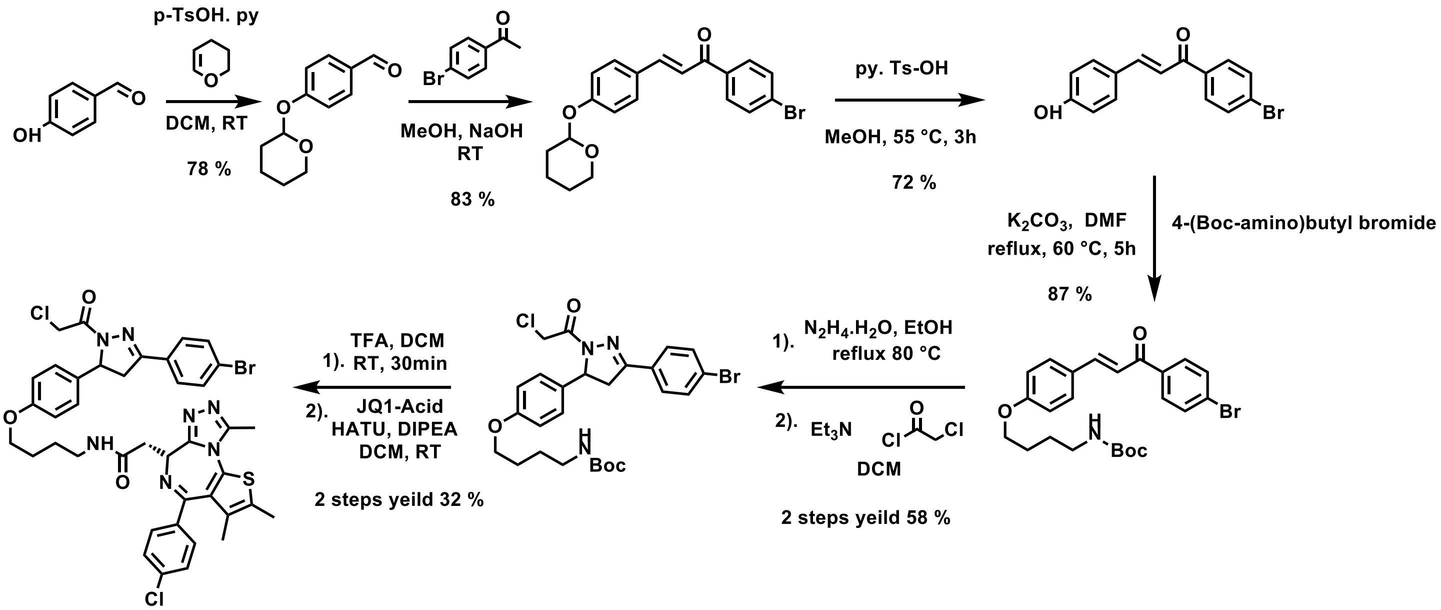


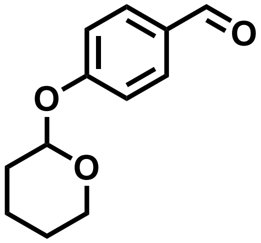

#### 4-((tetrahydro-2H-pyran-2-yl)oxy)benzaldehyde (ML 2-1)

To a solution of 4-hydroxybenzaldehyde (Alfa Aesar, 1.22 g, 10 mmol) in anhydrous DCM 3,4-Dihydro-2H-pyran (Sigma, 2.1 g, 25 mmol) and Pyridinium p-toluenesulfonate (Sigma, 0.122 g, 0.5 mmol) were added and the reaction stirred at room temperature overnight. Upon reaction completion the crude was purified by silica gel chromatography (20% EtOAc/hexanes) to yield 1.6 g (78%).
**$^1$H NMR (400 MHz $CDCl_3$):** δ 9.92 (s, 1H), 7.90 – 7.81 (m, 2H), 7.23 – 7.14 (m, 2H), 5.57 (t, *J* = 3.2 Hz, 1H), 3.87 (ddd, *J* = 11.3, 9.9, 3.1 Hz, 1H), 3.66 (dtd, *J* = 11.4, 4.1, 1.4 Hz, 1H), 2.12 – 1.98 (m, 1H), 1.98 – 1.83 (m, 2H), 1.82 – 1.54 (m, 3H).

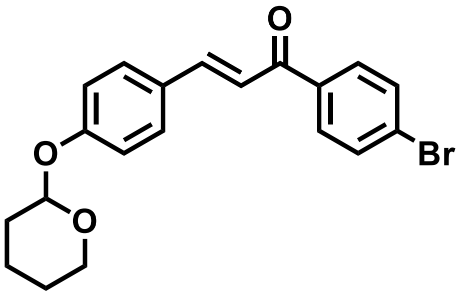

#### (E)-1-(4-bromophenyl)-3-(4-((tetrahydro-2H-pyran-2-yl)oxy)phenyl)prop-2-en-1-one (ML 2-2)

Equimolar portions of 4'-Bromacetophenone (Sigma, 1.45 g, 7.3 mmol) and ML 2-1 (1.5 g, 7.3 mmol) were dissolved in 20 mL ethanol, the reaction solution was stirred in ice bath for 15 min. 20 mL aliquot of 10% NaOH was then slowly added dropwise to the reaction solution. After stirring for 20 minutes, the reaction mixture was

allowed to warm to room temperature and was stirred for 3 hours. The white solid product was been filtered then washed with cold methanol and water, without further purification to yield 2.34 g (83%).
**$^1$H NMR (400 MHz $CDCl_3$):** δ 7.95 – 7.89 (m, 2H), 7.83 (d, *J* = 15.6 Hz, 1H), 7.71 – 7.61 (m, 4H), 7.41 (d, *J* = 15.6 Hz, 1H), 7.17 – 7.10 (m, 2H), 5.54 (t, *J* = 3.2 Hz, 1H), 3.92 (ddd, *J* = 11.4, 9.7, 3.1 Hz, 1H), 3.67 (dtd, *J* = 11.3, 4.1, 1.4 Hz, 1H), 2.15 – 2.00 (m, 1H), 1.92 (hd, *J* = 9.2, 8.3, 4.3 Hz, 2H), 1.81 – 1.66 (m, 3H).

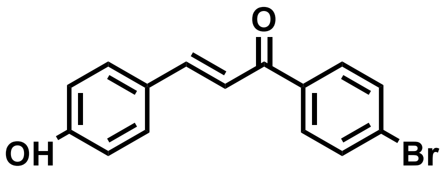


**(E)-1-(4-bromophenyl)-3-(4-hydroxyphenyl)prop-2-en-1-one (ML 2-3)**

To a solution of ML 2-2 (1.94 g, 5 mmol) in methanol (25 mL) Pyridinium p-toluenesulfonate (Sigma, 0.122 g, 0.5 mmol) were added. The reaction solution was stirred at 55˚ C for 3 h. Upon reaction completion the crude was purified by silica gel chromatography (30% EtOAc/hexanes) to yield 1.1 g (72%).
**$^1$H NMR (400 MHz, Methanol-$d_4$):** δ 7.98 (t, *J* = 10.8 Hz, 2H), 7.78 – 7.50 (m, 6H), 6.87 (d, *J* = 8.6 Hz, 2H).

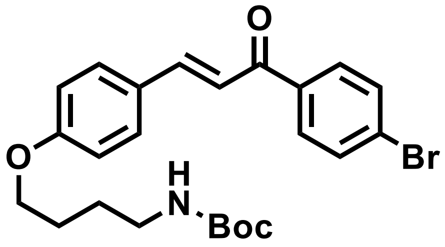


**(E)-tert-butyl (4-(4-(3-(4-bromophenyl)-3-oxoprop-1-en-1-yl)phenoxy)butyl)carbamate (ML 2-4)**

To a solution of ML 2-3 (226 mg, 0.75 mmol) in anhydrous DMF (8 mL), 4-(Boc-amino)butyl bromide (Sigma, 376 mg, 1.5 mmol) and $K_2CO_3$ (414 mg, 3 mmol) were added. The reaction solution was stirred at 60 ˚ C for 5 h under $N_2$ atmosphere. Upon reaction completion, the inorganic salts were filtered off, the solution was diluted with EtOAc, washed with water. The crude was purified by silica gel chromatography (25% EtOAc/hexanes) to yield 310 mg (87%).
**$^1$H NMR (400 MHz, $CDCl_3$):** δ 7.96 – 7.88 (m, 2H), 7.83 (d, *J* = 15.6 Hz, 1H), 7.72 – 7.59 (m, 4H), 7.39 (d, *J* = 15.6 Hz, 1H), 7.01 – 6.91 (m, 2H), 4.66 (s, 1H), 4.07 (t, *J* = 6.2 Hz, 2H), 3.25 (q, *J* = 6.8 Hz, 2H), 1.94 – 1.82 (m, 2H), 1.79 – 1.64 (m, 3H), 1.49 (s, 9H).

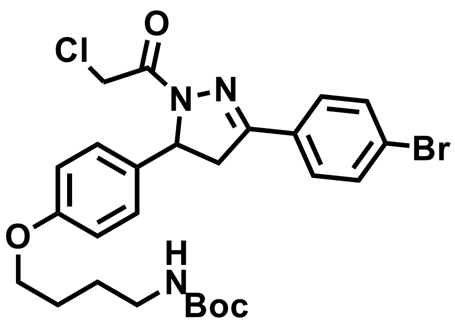


**tert-butyl (4-(4-(3-(4-bromophenyl)-1-(2-chloroacetyl)-4,5-dihydro-1H-pyrazol-5-yl)phenoxy)butyl)carbamate (ML 2-6)**

To a solution of ML 2-4 (550 mg, 1.16 mmol) in EtOH, hydrazine monohydrate (Sigma, 116 mg, 2.32 mmol) was added. The reaction solution was stirred at 80 ˚ C for 5 h under $N_2$ atmosphere. Upon reaction completion, the solution has diluted by water and extracted with DCM, combined the organic phase then dried it by anhydrous magnesium sulfate. The volume of the organic phase was evaporated *in vacuo* to around 5 mL. To the result solution, chloroacetyl chloride (Sigma, 157 mg, 1.39 mmol) and trimethylamine (152 mg, 1.5 mmol) were added. The reaction solution was stirred in ice bath for 30 min, then the reaction mixture was allowed to warm to room temperature and was stirred for overnight under $N_2$ atmosphere. Upon reaction completion, the reaction solution was diluted with EtOAc, washed with brine. The crude was purified by silica gel chromatography (40% EtOAc/hexanes) to yield 381 mg (58%).
**$^1$H NMR (400 MHz, $CDCl_3$):** δ 7.60 (q, *J* = 8.8 Hz, 4H), 7.19 – 7.10 (m, 2H), 6.87 – 6.77 (m, 2H), 5.55 (dd, *J* = 11.7, 4.7 Hz, 1H), 4.55 (s, 2H), 3.94 (t, *J* = 6.2 Hz, 2H), 3.74 (dd, *J* = 17.8, 11.7 Hz, 1H), 3.24 – 3.11 (m, 3H), 1.85 – 1.75 (m, 2H), 1.65 (p, *J* = 7.1 Hz, 2H), 1.44 (s, 9H).

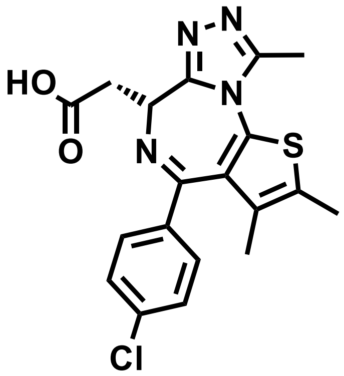

**(S)-2-(4-(4-chlorophenyl)-2,3,9-trimethyl-6H-thieno[3,2-f][1,2,4]triazolo[4,3-a][1,4]diazepin-6-yl)acetic acid (JQ1-acid)**

tert-butyl (S)-2-(4-(4-chlorophenyl)-2,3,9-trimethyl-6H-thieno[3,2-f][1,2,4]triazolo[4,3-a][1,4]diazepin-6-yl)acetate (JQ1, eNovation Chemicals, 204 mg, 0.446 mmol) was dissolved in formic acid (3mL) and stirred at 45 °C overnight. The mixture was diluted with DCM and solvent removed in vacuo. The resulting yellow oil was redissolved in 3 mL DCM and evaporated to dryness repeated until the process gave a fine yellow-brown solid: 178 mg (99.8%). No further purification was necessary.
**$^1$H NMR (400 MHz, $CDCl_3$):** δ 7.46 – 7.40 (m, 2H), 7.34 (d, J = 8.7 Hz, 2H), 4.59 (t, J = 6.8 Hz, 1H), 3.75 – 3.55 (m, 2H), 2.68 (s, 3H), 2.41 (s, 3H), 1.74 – 1.65 (m, 3H).

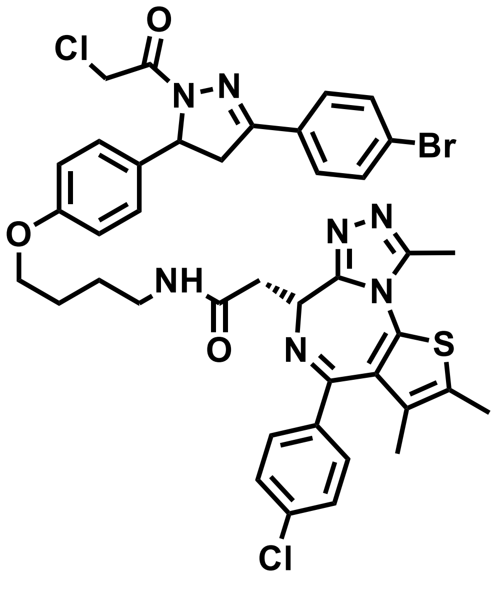

**N-(4-(4-(3-(4-bromophenyl)-1-(2-chloroacetyl)-4,5-dihydro-1H-pyrazol-5-yl)phenoxy)butyl)-2-((6R)-4-(4-chlorophenyl)-2,3,9-trimethyl-6H-thieno[3,2-f][1,2,4]triazolo[4,3-a][1,4]diazepin-6-yl)acetamide (ML 2-14)**

ML 2-6 (185 mg, 0.327 mmol) was deprotected in 1:1 DCM/TFA (5mL) by adding trifluoroacetic acid (SigmaAldrich) slowly over 20 min followed by stirring for an additional 20 min. TLC showed full conversion to amine and the solvent was removed in *vacuo,* and chases three times with 3mL DCM to remove excess TFA. The deprotected crude was used without further purification for amide coupling.
The resulting TFA salt was dissolved in 8 mL DCM, JQ1-acid (157 mg, 0.4 mmol), 1-[Bis(dimethylamino)methylene]-1H-1,2,3-triazolo[4,5-b]pyridinium 3-oxid hexafluorophosphate (HATU) (Small Molecules Inc., 202 mg, 0.530 mmol), and N,N-Diisopropylethylamine (DIPEA) (Sigma-Aldrich, 168 mg, 1.31 mmol). Stirred overnight and monitored by TLC (5% MeOH in DCM, 100% EtOAc). Crude concentrated and applied direct to a silica column for flash chromatography (1-5% MeOH/DCM). The eluted fractions were insufficiently pure and those containing product were combined, concentrated and purified again by flash silica chromatography (100-0% EtOAc/DCM followed by 0-5% MeOH/DCM) to afford 88.4 mg (32%)
**$^{1}$H NMR (400 MHz, $CDCl_3$):** δ 7.66 – 7.46 (m, 4H), 7.39 (d, *J* = 8.2 Hz, 2H), 7.29 (d, *J* = 8.3 Hz, 2H), 7.13 (d, *J* = 8.3 Hz, 2H), 6.86 (t, *J* = 5.9 Hz, 1H), 6.80 (d, *J* = 8.3 Hz, 2H), 5.54 (dt, *J* = 11.7, 4.1 Hz, 1H), 4.63 (t, *J* = 7.0 Hz, 1H), 4.54 (s, 2H), 3.90 (t, *J* = 6.1 Hz, 2H), 3.72 (dd, *J* = 17.8, 11.7 Hz, 1H), 3.54 (dd, *J* = 14.3, 7.5 Hz, 1H), 3.33 (ddt, *J* = 30.5, 13.3, 6.8 Hz, 3H), 3.17 (dd, *J* = 17.9, 4.7 Hz, 1H), 2.64 (s, 3H), 2.39 (s, 3H), 1.84 – 1.72 (m, 2H), 1.69 (d, *J* = 7.4 Hz, 2H), 1.65 (s, 3H).
**$^{13}$C NMR (151 MHz, $CDCl_3$)** δ 169.63, 163.05, 160.38, 157.82, 154.81, 153.53, 150.46, 149.02, 135.94, 135.73, 131.92, 131.27, 131.22, 130.05, 130.00, 129.61, 129.00, 128.95, 127.86, 127.37, 126.23, 124.29, 119.75, 114.06, 66.59, 59.36, 53.70, 41.30, 41.16, 38.67, 38.39, 25.72, 25.40, 13.52, 13.27, 12.24, 10.97, 10.58.
**HRMS (+ESI):** calcd. $C_{40}H_{39}BrCl_2N_7O_3S$ for = 846.1390; found 846.1399.

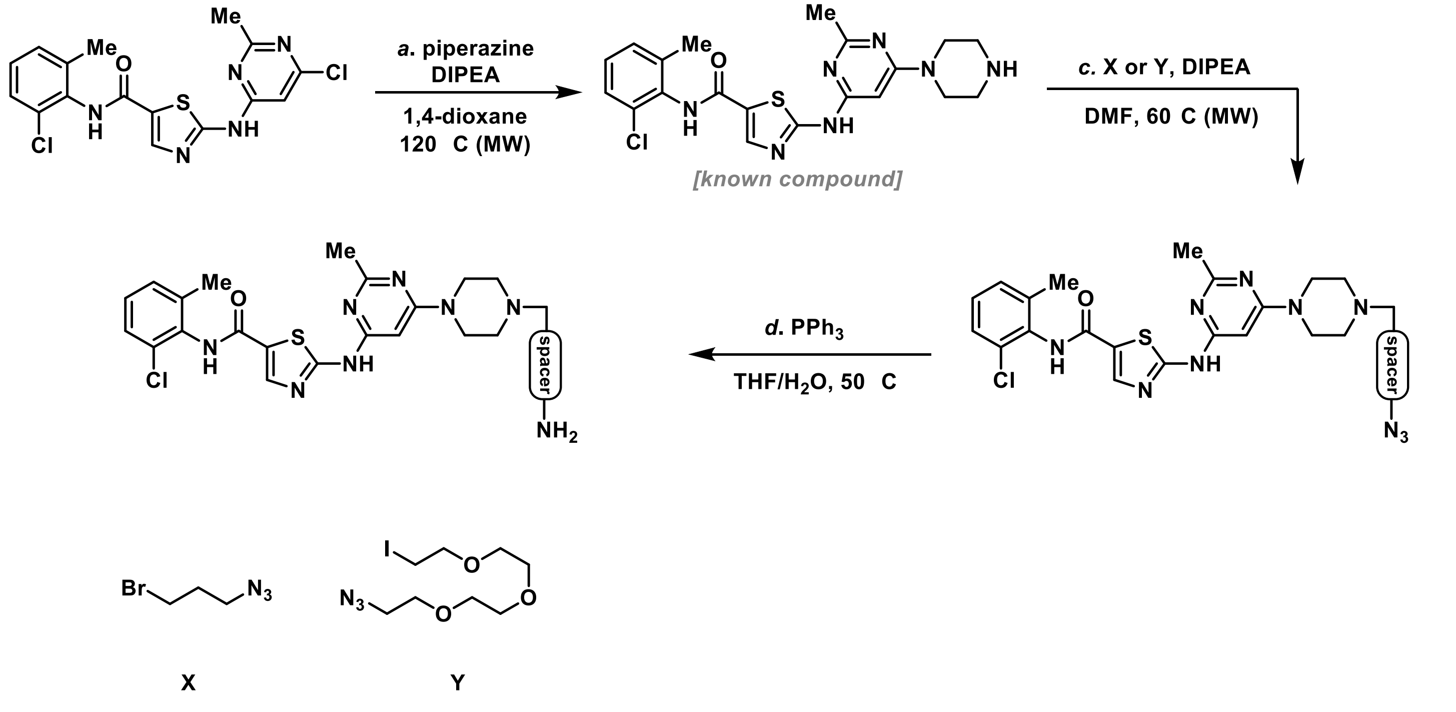

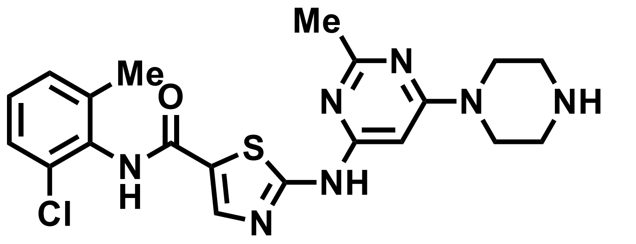

**Compound A**: This compound was synthesized according to the procedures reported by Veach, D. R. and coworkers. (*J. Med. Chem.* **2007**, *50*, 5853-5857)

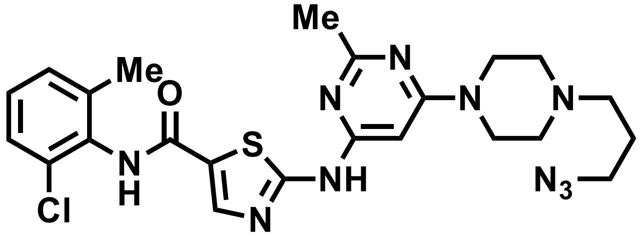

**Azide B**: A 30mL microwave tube equipped with a magnetic stir bar was charged with compound A (533 mg, 1.20 mmol, 1 equiv), DMF (12 mL), DIPEA (0.418 mL, 2.4 mmol, 2 equiv) and bromide X (0.178 mL, 1.44 mmol, 1.2 equiv). The tube was sealed and placed in the microwave reactor (Biotage®). The microwave reaction was performed at 60 °C for 1.5 h. After cooling to room temperature, the reaction mixture was transferred into a 100 mL round-bottom flask and distilled *in vacuo* in order to remove DMF and other volatiles. The remaining white solid was then suspended in DCM/MeOH (90:10 v/v), passed through a short column of silica gel and eluted with DCM/MeOH (90:10 v/v). The resulting solution was concentrated *in vacuo* to afford the crude product which was further purified by column chromatography (DCM:MeOH = 95:5) to yield azide B (290 mg, 46% yield) as a white solid.
**$^{1}$H NMR (600 MHz, $(CD_3)_2SO$)** δ 11.47 (s, 1H), 9.88 (s, 1H), 8.22 (s, 1H), 7.40 (d, *J* = 7.7 Hz, 1H), 7.33 – 7.22 (m, 2H), 6.05 (s, 1H), 3.51 (br, 4H), 3.39 (t, *J* = 6.7 Hz, 2H), 2.43 (br, 4H), 2.40 (s, 3H), 2.38 (t, *J* = 6.9 Hz, 2H), 2.24 (s, 3H), 1.73 (p, *J* = 6.8 Hz, 2H).
**$^{13}$C NMR (151 MHz, $(CD_3)_2SO$)** δ 165.2, 162.6, 162.4, 159.9, 156.9, 140.8, 138.8, 133.5, 132.4, 129.0, 128.1, 127.0, 125.7, 82.6, 54.6, 52.2, 48.9, 43.6, 25.61, 25.58, 18.3.
**IR (thin film, cm$^{-1}$)** 3205, 3060, 2938, 2847, 2093, 1715, 1621, 1536, 1182, 1142, 998, 862. **HRMS (ESI)** *calcd* for $[C_{23}H_{28}ON_{10}ClS]^+$ ($[M+H]^+$): *m/z* 527.1851, found: 527.1850.

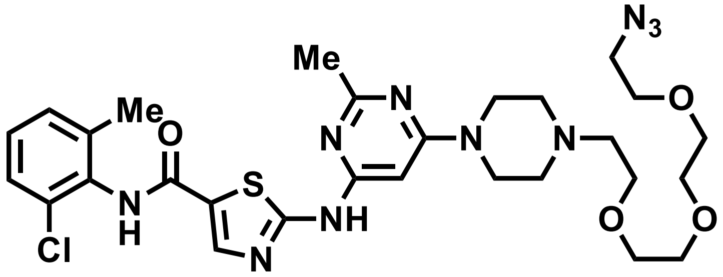

**Azide C**: A 5 mL microwave tube equipped with a magnetic stir bar was charged with compound A (111 mg, 0.25 mmol, 1 equiv), DMF (1.5 mL), DIPEA (65.0 µL, 0.375 mmol, 1.5 equiv) and iodide Y (51 µL, 0.25 mmol, 1.0 equiv). The tube was sealed and placed in the microwave reactor. The microwave reaction was performed at 60 °C for 1 h. After cooling to room temperature, the reaction mixture was transferred into a 50 mL round-bottom flask and the solvent and volatiles were remove under high vacuum. The remaining white solid was

purified by column chromatography (DCM:MeOH = 95:5 to 93:7) to afford azide C (101 mg, 63% yield) as a white solid.
**$^1$H NMR (600 MHz, MeOD)** δ 8.15 (s, 1H), 7.35 (d, *J* = 7.2 Hz, 1H), 7.30 – 7.18 (m, 2H), 6.02 (s, 1H), 3.72 – 3.62 (m, 16H), 3.37 (t, *J* = 4.8 Hz, 2H), 2.70 (t, *J* = 5.4 Hz, 2H), 2.68 (br, 4H), 2.48 (s, 3H), 2.32 (s, 3H).
**$^{13}$C NMR (151 MHz, MeOD)** δ 167.5, 165.3, 164.5, 163.3, 158.6, 142.2, 140.4, 134.4, 134.3, 130.1, 129.5, 128.3, 126.8, 83.9, 71.7, 71.7, 71.6, 71.4, 71.2, 69.4, 58.7, 54.1, 51.8, 44.7, 25.6, 18.7.
**IR (thin film, cm$^{-1}$)** 3182, 3061, 2925, 2870, 2094, 1718, 1616, 1536, 1189, 1144, 1010, 952, 781.
**HRMS (ESI)** *calcd* for $[C_{28}H_{38}O_4N_{10}ClS]^+$ ($[M+H]^+$) *m/z* 645.2481, found: 645.2476.

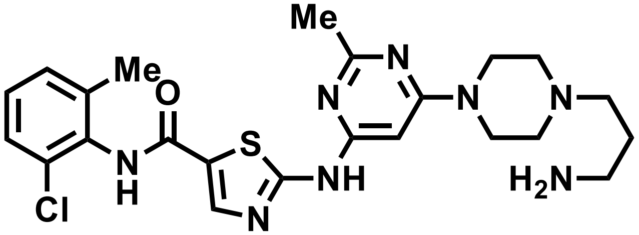


**Amine D**: To a 10 mL reaction tube was added azide B (20.8 mg, 0.0395 mmol, 1.0 equiv), triphenylphosphine (16 mg, 0.059 mmol, 1.5 equiv), THF (1.0 mL) and $H_2O$ (0.1 mL). The resulting solution was heated in a 50 °C oil bath for 12 hours. After fully consumption of the starting material was indicated by TLC, all the volatiles in the reaction mixture was removed by rotary evaporation at 60 °C. The resulting crude pale-yellow solid containing Amine D was further dried under high vacuum overnight before being subjected to the next step.

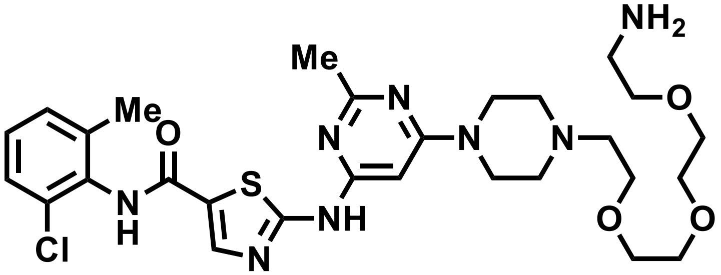


**Amine E**: The same procedure as the synthesis of Amine D was applied using azide C (11.7 mg, 0.0181 mmol, 1.0 equiv), triphenylphosphine (7.1 mg, 0.027 mmol, 1.5 equiv), THF (0.8 mL) and $H_2O$ (0.1 mL) to afford crude Amine E.

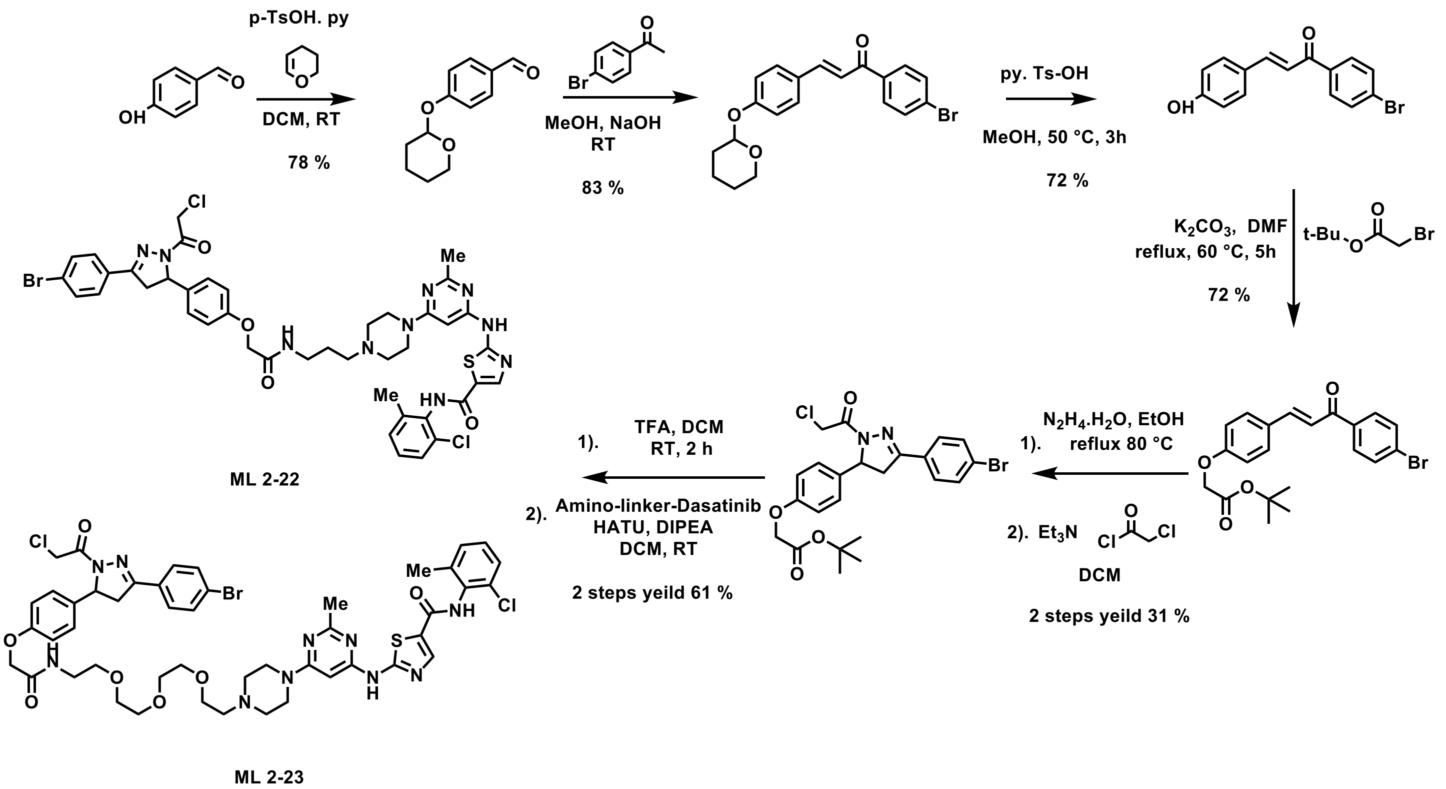


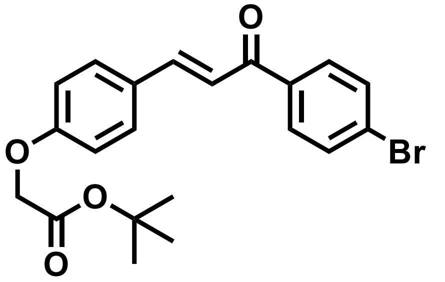

**(E)-tert-butyl 2-(4-(3-(4-bromophenyl)-3-oxoprop-1-en-1-yl)phenoxy)acetate (ML 2-20)**

To a solution of ML 2-3 (153 mg, 0.50 mmol) in anhydrous DMF (8 mL), tert-Butyl bromoacetate (Sigma, 146 mg, 0.75 mmol) and $K_2CO_3$ (138 mg, 1 mmol) were added. The reaction solution was stirred at 60 ˚ C for 5 h under $N_2$ atmosphere. Upon reaction completion, the inorganic salts were filtered off, the solution was diluted with EtOAc, washed with water. The crude was purified by silica gel chromatography (25% EtOAc/hexanes) to yield 150 mg (72%).
**$^1$H NMR (400 MHz, $CDCl_3$):** δ 7.97 – 7.89 (m, 2H), 7.82 (d, *J* = 15.6 Hz, 1H), 7.73 – 7.60 (m, 4H), 7.41 (d, *J* = 15.6 Hz, 1H), 7.01 – 6.93 (m, 2H), 4.61 (s, 2H), 1.54 (s, 9H).

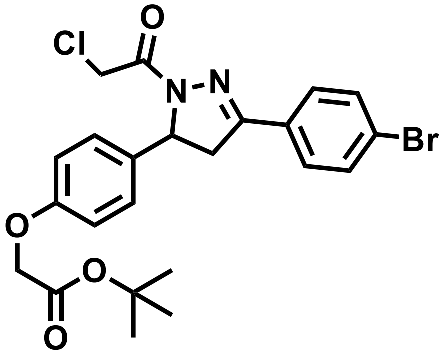

***tert*-butyl 2-(4-(3-(4-bromophenyl)-1-(2-chloroacetyl)-4,5-dihydro-1H-pyrazol-5-yl)phenoxy)acetate (ML 2-21)**

To a solution of ML 2-20 (417 mg, 1.00 mmol) in EtOH, hydrazine monohydrate (Sigma, 250 mg, 5.00 mmol) was added. The reaction solution was stirred at 80 ˚ C for 5 h under $N_2$ atmosphere. Upon reaction completion, the solution has diluted by water and extracted with DCM, combined the organic phase then dried it by anhydrous magnesium sulfate. The volume of the organic phase was evaporated *in vacuo* to around 5 mL. To the result solution, chloroacetyl chloride (Sigma, 169 mg, 1.50 mmol) and trimethylamine (303 mg, 3.00 mmol) were added. The reaction solution was stirred in ice bath for 30 min, then the reaction mixture was allowed to warm to room temperature and was stirred for overnight under $N_2$ atmosphere. Upon reaction completion, the reaction solution was diluted with EtOAc, washed with brine. The crude was purified by silica gel chromatography (35 – 40 % EtOAc/hexanes) to yield 140 mg (28%).
**$^1$H NMR (400 MHz, $CDCl_3$):** δ 7.60 – 7.54 (m, 2H), 7.25 – 7.17 (m, 2H), 7.02 – 6.94 (m, 2H), 6.92 – 6.84 (m, 2H), 6.77 (s, 1H), 5.60 (dd, *J* = 11.7, 4.6 Hz, 1H), 4.60 (s, 2H), 4.51 (s, 2H), 3.77 (dd, *J* = 17.8, 11.7 Hz, 1H), 3.22 (dd, *J* = 17.8, 4.6 Hz, 1H), 1.54 (s, 9H).

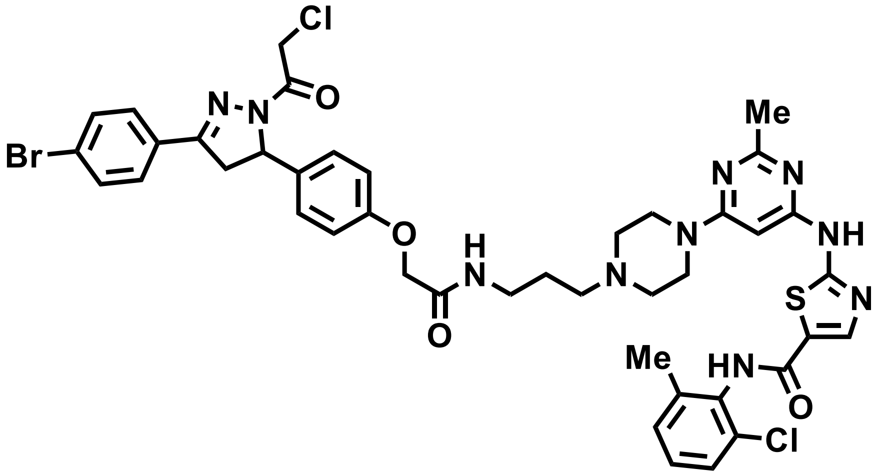


**2-((6-(4-(3-(2-(4-(3-(4-bromophenyl)-1-(2-chloroacetyl)-4,5-dihydro-1H-pyrazol-5-yl)phenoxy)acetamido)propyl)piperazin-1-yl)-2-methylpyrimidin-4-yl)amino)-N-(2-chloro-6-methylphenyl)thiazole-5-carboxamide (ML 2-22)**

ML 2-21 (15.2 mg, 0.030 mmol) was deprotected in 3:1 DCM/TFA (2 mL) by adding trifluoroacetic acid (SigmaAldrich) slowly over 20 min followed by stirring for an additional 20 min. TLC showed full conversion to amine and the solvent was removed in *vacuo,* and chases three times with 3mL DCM to remove excess TFA. The deprotected crude was used without further purification for amide coupling.
The resulting TFA salt was dissolved in 2 mL DCM, **Amine D** (15.0 mg, 0.030 mmol), 1-[Bis(dimethylamino)methylene]-1H-1,2,3-triazolo[4,5-b]pyridinium 3-oxid hexafluorophosphate (HATU) (Small Molecules Inc., 17.1 mg, 0.045 mmol), and N,N-Diisopropylethylamine (DIPEA) (Sigma-Aldrich, 193.5 mg, 1.50 mmol). Stirred overnight and monitored by TLC (20% Methanol in DCM). Upon reaction completion, the reaction solution was diluted with EtOAc, washed with brine. ). Crude concentrated and applied direct to a silica column for flash chromatography (1-5% MeOH/DCM). The eluted fractions were insufficiently pure and those containing product were combined, concentrated and purified again by flash silica chromatography (100-0% EtOAc/DCM followed by 10-35% MeOH/DCM) to afford 17.1 mg (60%)
**$^1$H NMR (600 MHz, DMSO-d6)** δ 11.47 (s, 1H), 9.88 (s, 1H), 8.23 (s, 1H), 8.11 (d, J = 18.3 Hz, 1H), 7.84 – 7.71 (m, 2H), 7.71 – 7.64 (m, 2H), 7.40 (dd, J = 7.8, 1.6 Hz, 1H), 7.34 – 7.22 (m, 2H), 7.22 – 7.09 (m, 2H), 6.99 – 6.85 (m, 2H), 6.06 (s, 1H), 5.54 (dd, J = 11.7, 4.8 Hz, 1H), 4.79 – 4.59 (m, 2H), 4.46 (s, 2H), 3.93 – 3.79 (m, 1H), 3.51 (d, J = 9.8 Hz, 4H), 3.26 – 3.07 (m, 3H), 2.41 (s, 6H), 2.31 (s, 2H), 2.25 (s, 3H), 1.67 (d, J = 40.6 Hz, 2H).
**$^{13}$C NMR (151 MHz, DMSO)** δ 166.99, 164.73, 162.75, 162.11, 161.94, 159.47, 156.62, 156.51, 154.33, 140.38, 138.38, 133.81, 133.08, 132.00, 131.36, 129.50, 129.04, 128.58, 128.38, 127.72, 126.54, 125.28,

123.65, 114.43, 82.24, 66.71, 59.24, 55.10, 54.47, 51.85, 43.14, 41.99, 41.37, 36.54, 29.48, 27.95, 25.13, 22.97, 21.95, 17.87, 13.49, 10.47.
**HRMS (+ESI):** calcd. $C_{42}H_{43}BrCl_2N_{10}O_4S$ for = 933.1823; found 933.1814

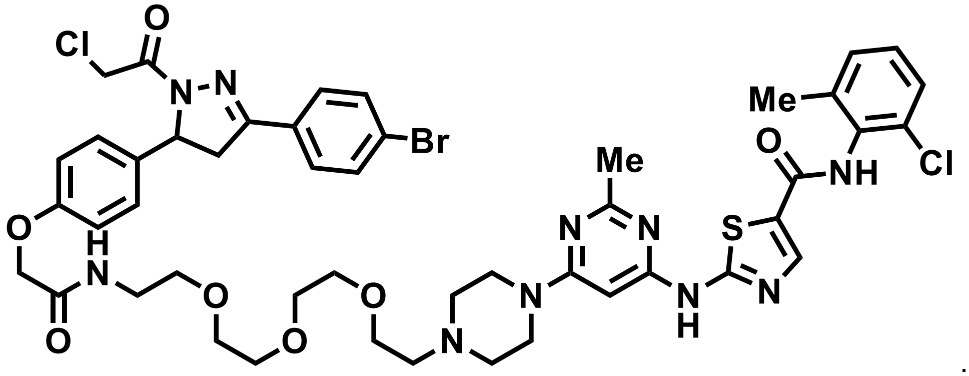


**2-((6-(4-(1-(4-(3-(4-bromophenyl)-1-(2-chloroacetyl)-4,5-dihydro-1H-pyrazol-5-yl)phenoxy)-2-oxo-6,9,12-trioxa-3-azatetradecan-14-yl)piperazin-1-yl)-2-methylpyrimidin-4-yl)amino)-N-(2-chloro-6-methylphenyl)thiazole-5-carboxamide (ML 2-23)**

ML 2-21 (16.2 mg, 0.032 mmol) was deprotected in 3:1 DCM/TFA (2 mL) by adding trifluoroacetic acid (SigmaAldrich) slowly over 20 min followed by stirring for an additional 20 min. TLC showed full conversion to amine and the solvent was removed in *vacuo,* and chases three times with 3mL DCM to remove excess TFA. The deprotected crude was used without further purification for amide coupling.
The resulting TFA salt was dissolved in 2 mL DCM, **Amine E** (19.8 mg, 0.032 mmol), 1-[Bis(dimethylamino)methylene]-1H-1,2,3-triazolo[4,5-b]pyridinium 3-oxid hexafluorophosphate (HATU) (Small Molecules Inc., 18.2 mg, 0.048 mmol), and N,N-Diisopropylethylamine (DIPEA) (Sigma-Aldrich, 206.0 mg, 1.60 mmol). Stirred overnight and monitored by TLC (20% Methanol in DCM). Upon reaction completion, the reaction solution was diluted with EtOAc, washed with brine. ). Crude concentrated and applied direct to a silica column for flash chromatography (1-5% MeOH/DCM). The eluted fractions were insufficiently pure and those containing product were combined, concentrated and purified again by flash silica chromatography (100-0% EtOAc/DCM followed by 10-35% MeOH/DCM) to afford 20.1 mg (59%)
**$^1$H NMR (400 MHz, Methanol-*d*$_4$)** δ 8.16 (s, 1H), 7.77 – 7.71 (m, 2H), 7.61 (dd, *J* = 8.8, 2.3 Hz, 2H), 7.40 – 7.34 (m, 1H), 7.31 – 7.15 (m, 4H), 7.00 – 6.90 (m, 2H), 6.00 (s, 1H), 5.57 (dd, *J* = 11.7, 4.7 Hz, 1H), 4.73 (d, *J* = 13.8 Hz, 1H), 4.60 (d, *J* = 13.9 Hz, 1H), 4.52 (s, 2H), 4.12 (q, *J* = 7.1 Hz, 1H), 3.88 (dd, *J* = 18.2, 11.7 Hz, 1H), 3.62 (tdd, *J* = 13.9, 10.2, 6.7 Hz, 16H), 3.48 (t, *J* = 5.5 Hz, 2H), 3.19 (dd, *J* = 18.1, 4.8 Hz, 1H), 2.66 (dd, *J* = 11.8, 5.7 Hz, 6H), 2.47 (s, 3H), 2.34 (s, 3H).
**$^{13}$C NMR (151 MHz, DMSO)** δ 167.22, 164.61, 162.76, 161.87, 159.47, 156.62, 154.36, 140.39, 137.64, 133.32, 133.08, 132.00, 131.38, 131.33, 129.05, 128.59, 128.40, 127.28, 126.54, 114.44, 69.29, 69.15, 68.36, 66.70, 59.25, 53.60, 42.00, 41.39, 41.27, 37.81, 35.34, 30.25, 29.48, 27.42, 25.11, 22.97, 21.95, 17.86, 17.60, 15.59, 13.46, 11.90, 10.47.
**HRMS (+ESI):** calcd. $C_{47}H_{54}BrCl_2N_{10}O_4S$ for = 1051.2453; found 1051.2445

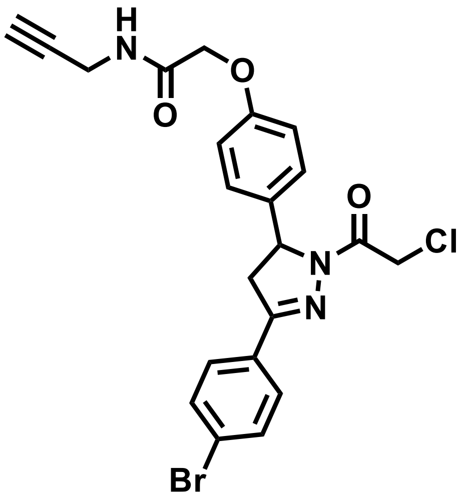

**2-(4-(3-(4-bromophenyl)-1-(2-chloroacetyl)-4,5-dihydro-1H-pyrazol-5-yl)phenoxy)-N-(prop-2-yn-1-yl)acetamide (ML 2-24)**

ML 2-21 (17.6 mg, 0.035 mmol) was deprotected in 3:1 DCM/TFA (2 mL) by adding trifluoroacetic acid (SigmaAldrich) slowly over 20 min followed by stirring for an additional 20 min. TLC showed full conversion to amine and the solvent was removed in *vacuo,* and chases three times with 3mL DCM to remove excess TFA. The deprotected crude was used without further purification for amide coupling.
The resulting TFA salt was dissolved in 2 mL DCM, propargylamine (2.3 mg, 0.042 mmol), 1-[Bis(dimethylamino)methylene]-1H-1,2,3-triazolo[4,5-b]pyridinium 3-oxid hexafluorophosphate (HATU) (Small Molecules Inc., 19.8 mg, 0.052 mmol), and N,N-Diisopropylethylamine (DIPEA) (Sigma-Aldrich, 223.9 mg, 1.750 mmol). Stirred overnight and monitored by TLC (60% EtOAc in hexane). Upon reaction completion, the reaction solution was diluted with DCM, washed with brine. The crude was purified by silica gel chromatography (25 – 70 % EtOAc/hexanes) to yield 14.6 mg (88%).
**$^1$H NMR (400 MHz, $CDCl_3$):** δ 7.68 – 7.60 (m, 4H), 7.27 – 7.21 (m, 2H), 6.95 – 6.89 (m, 2H), 6.81 (s, 1H), 5.60 (dd, *J* = 11.8, 4.8 Hz, 1H), 4.64 – 4.54 (m, 2H), 4.52 (s, 2H), 4.18 (dd, *J* = 5.6, 2.6 Hz, 2H), 3.81 (dd, *J* = 17.8, 11.8 Hz, 1H), 3.22 (dd, *J* = 17.9, 4.8 Hz, 1H), 2.31 (t, *J* = 2.6 Hz, 1H).
**$^{13}$C NMR (101 MHz, $CDCl_3$):** δ 167.87, 164.14, 156.91, 154.49, 134.70, 132.26, 128.35, 127.59, 125.46, 115.32, 77.48, 77.16, 76.84, 72.06, 67.44, 60.19, 42.24, 42.12, 28.86.
**HRMS (+ESI):** calcd. $C_{22}H_{19}BrCl_1N_3O_3Na$ for = 510.0191; found 510.0186

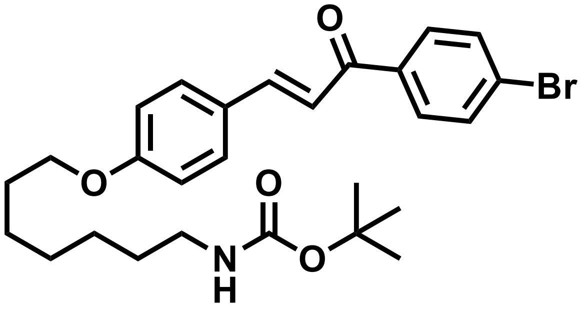

**(E)-*tert*-butyl (7-(4-(3-(4-bromophenyl)-3-oxoprop-1-en-1-yl)phenoxy)heptyl)carbamate (ML 2-27)**

To a solution of ML 2-3 (153 mg, 0.50 mmol) in anhydrous DMF (8 mL), N-Boc-7-bromoheptan-1-amine (AChemBlock, 300 mg, 1.00 mmol) and $K_2CO_3$ (276 mg, 2.00 mmol) were added. The reaction solution was stirred at 60 ˚ C for 5 h under $N_2$ atmosphere. Upon reaction completion, the inorganic salts were filtered off, the solution was diluted with EtOAc, washed with water. The crude was purified by silica gel chromatography (25% EtOAc/hexanes) to yield 238 mg (92%).

**$^{1}$H NMR (400 MHz, CDCl$_3$):** δ 7.97 – 7.89 (m, 2H), 7.83 (d, *J* = 15.6 Hz, 1H), 7.73 – 7.59 (m, 4H), 7.39 (d, *J* = 15.6 Hz, 1H), 7.00 – 6.92 (m, 2H), 4.55 (s, 1H), 4.04 (t, *J* = 6.5 Hz, 2H), 3.16 (q, *J* = 6.8 Hz, 2H), 1.94 – 1.76 (m, 2H), 1.57 – 1.50 (m, 4H), 1.49 (s, 9H), 1.44 – 1.37 (m, 4H).

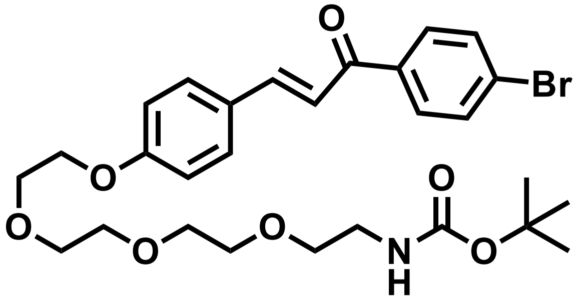


**(E)-tert-butyl (2-(2-(2-(2-(4-(3-(4-bromophenyl)-3-oxoprop-1-en-1-yl)phenoxy)ethoxy)ethoxy)ethoxy)ethyl)carbamate (ML 2-28)**

To a solution of ML 2-3 (153 mg, 0.50 mmol) in anhydrous DMF (8 mL), tert-Butyl (2-(2-(2-(2-bromoethoxy)ethoxy)ethoxy)ethyl)carbamate (Synthonix, 344 mg, 0.97 mmol) and $K_2CO_3$ (276 mg, 2.00 mmol) were added. The reaction solution was stirred at 60 ˚ C for 5 h under $N_2$ atmosphere. Upon reaction completion, the inorganic salts were filtered off, the solution was diluted with EtOAc, washed with water. The crude was purified by silica gel chromatography (50% EtOAc/hexanes) to yield 251 mg (88%).
**$^{1}$H NMR (400 MHz, CDCl$_3$):** δ 7.97 – 7.89 (m, 2H), 7.89 – 7.78 (m, 1H), 7.72 – 7.55 (m, 4H), 7.40 (d, *J* = 15.6 Hz, 1H), 7.04 – 6.95 (m, 2H), 5.07 (s, 1H), 4.23 (dd, *J* = 5.7, 4.0 Hz, 2H), 3.97 – 3.84 (m, 2H), 3.83 – 3.77 (m, 2H), 3.76 – 3.72 (m, 2H), 3.71 – 3.64 (m, 4H), 3.58 (t, *J* = 5.1 Hz, 2H), 3.35 (d, *J* = 6.3 Hz, 2H), 1.48 (s, 9H).

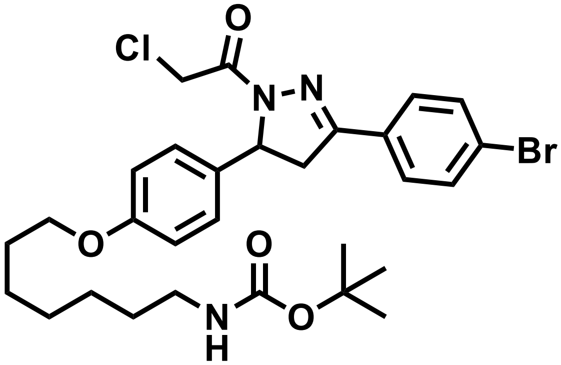


***tert*-butyl (7-(4-(3-(4-bromophenyl)-1-(2-chloroacetyl)-4,5-dihydro-1H-pyrazol-5-yl)phenoxy)heptyl)carbamate (ML 2-29)**

To a solution of ML 2-27 (223 mg, 0.43 mmol) in EtOH, hydrazine monohydrate (Sigma, 43 mg, 0.86 mmol) was added. The reaction solution was stirred at 80 ˚ C for 3 h under $N_2$ atmosphere. Upon reaction completion, the solution has diluted by water and extracted with DCM, combined the organic phase then dried it by anhydrous magnesium sulfate. The volume of the organic phase was evaporated *in vacuo* to around 5 mL. To the result solution, chloroacetyl chloride (Sigma, 58 mg, 0.52 mmol) and trimethylamine (57 mg, 0.56 mmol) were added. The reaction solution was stirred in ice bath for 30 min, then the reaction mixture was allowed to warm to room temperature and was stirred for overnight under $N_2$ atmosphere. Upon reaction completion, the reaction solution was diluted with EtOAc, washed with brine. The crude was purified by silica gel chromatography (25 – 40 % EtOAc/hexanes) to yield 104 mg (40%).
**$^{1}$H NMR (400 MHz, CDCl$_3$):** δ 7.69 – 7.53 (m, 4H), 7.23 – 7.12 (m, 2H), 6.89 – 6.82 (m, 2H), 5.57 (dd, *J* = 11.7, 4.7 Hz, 1H), 4.57 (s, 2H), 3.93 (t, *J* = 6.5 Hz, 2H), 3.76 (dd, *J* = 17.8, 11.7 Hz, 1H), 3.21 (dd, *J* = 17.9, 4.7 Hz, 1H), 3.13 (q, *J* = 6.8 Hz, 2H), 1.85 – 1.70 (m, 2H), 1.55 – 1.46 (m, 4H), 1.47 (s, 9H), 1.41 – 1.32 (m, 4H).

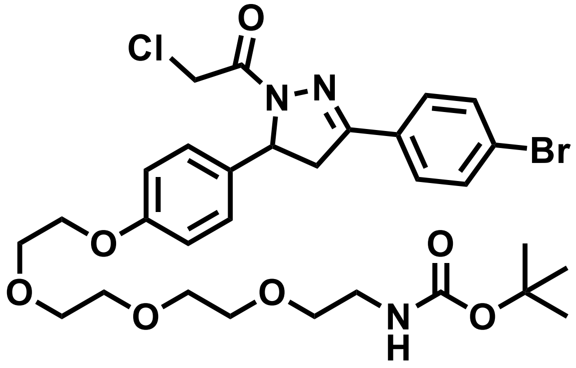

***tert*-butyl (2-(2-(2-(2-(4-(3-(4-bromophenyl)-1-(2-chloroacetyl)-4,5-dihydro-1H-pyrazol-5-yl)phenoxy)ethoxy)ethoxy)ethoxy)ethyl)carbamate (ML 2-30)**

To a solution of ML 2-28 (244 mg, 0.42 mmol) in EtOH, hydrazine monohydrate (Sigma, 42 mg, 0.84 mmol) was added. The reaction solution was stirred at 80 ˚ C for 5 h under $N_2$ atmosphere. Upon reaction completion, the solution has diluted by water and extracted with DCM, combined the organic phase then dried it by anhydrous magnesium sulfate. The volume of the organic phase was evaporated *in vacuo* to around 5 mL. To the result solution, chloroacetyl chloride (Sigma, 58 mg, 0.52 mmol) and trimethylamine (57 mg, 0.56 mmol) were added. The reaction solution was stirred in ice bath for 30 min, then the reaction mixture was allowed to warm to room temperature and was stirred for overnight under $N_2$ atmosphere. Upon reaction completion, the reaction solution was diluted with EtOAc, washed with brine. The crude was purified by silica gel chromatography (30 – 65 % EtOAc/hexanes) to yield 82 mg (30%).
**$^1$H NMR (400 MHz, $CDCl_3$):** δ 7.67 – 7.58 (m, 4H), 7.21 – 7.14 (m, 2H), 6.92 – 6.85 (m, 2H), 5.57 (dd, *J* = 11.7, 4.7 Hz, 1H), 4.57 (d, *J* = 1.1 Hz, 2H), 4.12 (dd, *J* = 5.6, 4.2 Hz, 2H), 3.88 – 3.83 (m, 2H), 3.75 – 3.73 (m, 2H), 3.72 – 3.69 (m, 2H), 3.67 – 3.62 (m, 4H), 3.55 (t, *J* = 5.1 Hz, 2H), 3.32 (q, *J* = 4.3, 3.4 Hz, 2H), 3.21 (dd, *J* = 17.8, 4.7 Hz, 1H), 1.46 (s, 9H).

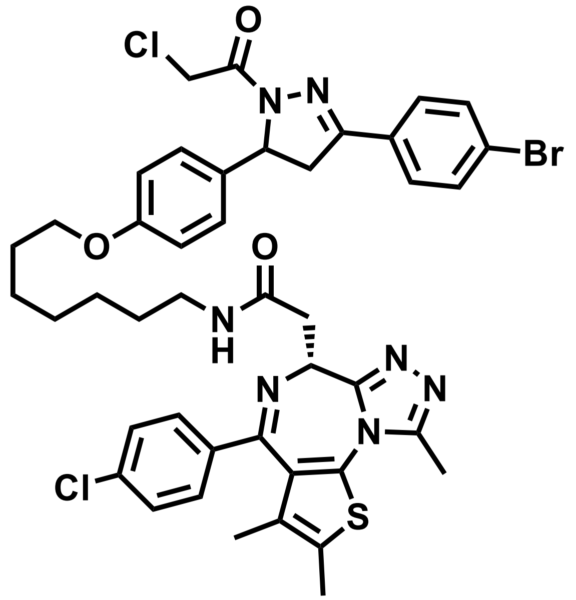

**N-(7-(4-(3-(4-bromophenyl)-1-(2-chloroacetyl)-4,5-dihydro-1H-pyrazol-5-yl)phenoxy)heptyl)-2-((6R)-4-(4-chlorophenyl)-2,3,9-trimethyl-6H-thieno[3,2-f][1,2,4]triazolo[4,3-a][1,4]diazepin-6-yl)acetamide (ML 2-31)**

ML 2-29 (104 mg, 0.171 mmol) was deprotected in 1:1 DCM/TFA (5mL) by adding trifluoroacetic acid (SigmaAldrich) slowly over 20 min followed by stirring for an additional 20 min. TLC showed full conversion to

amine and the solvent was removed in *vacuo,* and chases three times with 3mL DCM to remove excess TFA. The deprotected crude was used without further purification for amide coupling.
The resulting TFA salt was dissolved in 8 mL DCM, JQ1-acid (82 mg, 0.205 mmol), 1-[Bis(dimethylamino)methylene]-1H-1,2,3-triazolo[4,5-b]pyridinium 3-oxid hexafluorophosphate (HATU) (Small Molecules Inc., 97.5 mg, 0.257 mmol), and N,N-Diisopropylethylamine (DIPEA) (Sigma-Aldrich, 551.5 mg, 4.275 mmol). Stirred overnight and monitored by TLC (5% MeOH in DCM, 100% EtOAc). Crude concentrated and applied direct to a silica column for flash chromatography (1-5% MeOH/DCM). The eluted fractions were insufficiently pure and those containing product were combined, concentrated and purified again by flash silica chromatography (100-0% EtOAc/DCM followed by 0-5% MeOH/DCM) to afford 66 mg (43%)
**$^{1}$H NMR (400 MHz, $CDCl_3$):** δ 7.68 – 7.57 (m, 4H), 7.48 – 7.39 (m, 2H), 7.39 – 7.32 (m, 2H), 7.22 – 7.11 (m, 2H), 6.90 – 6.81 (m, 2H), 6.67 (t, *J* = 5.8 Hz, 1H), 5.58 (ddd, *J* = 11.8, 4.7, 2.1 Hz, 1H), 4.69 – 4.61 (m, 1H), 4.59 (d, *J* = 2.5 Hz, 2H), 3.93 (t, *J* = 6.4 Hz, 2H), 3.84 – 3.67 (m, 2H), 3.56 (dd, *J* = 14.3, 7.4 Hz, 1H), 3.42 – 3.30 (m, 2H), 3.28 – 3.16 (m, 3H), 2.70 (s, 3H), 2.47 – 2.35 (m, 3H), 1.82 – 1.71 (m, 2H), 1.71 – 1.67 (m, 3H), 1.56 (p, *J* = 7.0 Hz, 2H), 1.45 – 1.43 (m, 4H).
**$^{13}$C NMR (151 MHz, $CDCl_3$)** δ 170.39, 163.89, 158.85, 155.68, 154.57, 149.87, 136.78, 136.60, 132.63, 132.09, 132.06, 130.94, 130.89, 130.46, 129.83, 128.70, 128.24, 127.00, 125.15, 115.09, 115.05, 114.91, 67.94, 60.24, 54.91, 54.51, 42.98, 42.21, 42.06, 39.62, 39.37, 29.69, 29.43, 29.12, 28.99, 26.82, 25.94, 18.54, 17.15, 14.38, 13.08, 12.44, 11.80.
**HRMS (+ESI):** calcd. $C_{43}H_{45}BrCl_2N_7O_3S$ for = 888.1865; found 888.1858

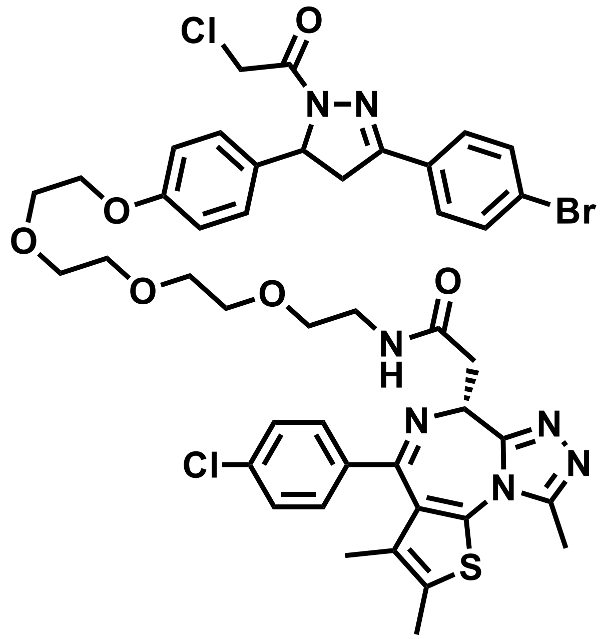


**N-(2-(2-(2-(2-(4-(3-(4-bromophenyl)-1-(2-chloroacetyl)-4,5-dihydro-1H-pyrazol-5-yl)phenoxy)ethoxy)ethoxy)ethoxy)ethyl)-2-((6R)-4-(4-chlorophenyl)-2,3,9-trimethyl-6H-thieno[3,2-f][1,2,4]triazolo[4,3-a][1,4]diazepin-6-yl)acetamide (ML 2-32)**

ML 2-30 (81.7 mg, 0.120 mmol) was deprotected in 1:1 DCM/TFA (5mL) by adding trifluoroacetic acid (SigmaAldrich) slowly over 20 min followed by stirring for an additional 20 min. TLC showed full conversion to amine and the solvent was removed in *vacuo,* and chases three times with 3mL DCM to remove excess TFA. The deprotected crude was used without further purification for amide coupling.
The resulting TFA salt was dissolved in 8 mL DCM, JQ1-acid (57.6 mg, 0.150 mmol), 1-[Bis(dimethylamino)methylene]-1H-1,2,3-triazolo[4,5-b]pyridinium 3-oxid hexafluorophosphate (HATU) (Small Molecules Inc., 68.4 mg, 0.180 mmol), and N,N-Diisopropylethylamine (DIPEA) (Sigma-Aldrich, 774 mg, 6.00 mmol). Stirred overnight and monitored by TLC (10% MeOH in DCM, 100% EtOAc). Crude concentrated and applied direct to a silica column for flash chromatography (1-10% MeOH/DCM). The eluted fractions were insufficiently pure and those containing product were combined, concentrated and purified again by flash silica chromatography (100-0% EtOAc/DCM followed by 0-10% MeOH/DCM) to afford 46 mg (40%)

**$^{1}$H NMR (400 MHz, $CDCl_3$):** δ 7.66 – 7.57 (m, 4H), 7.45 – 7.41 (m, 2H), 7.34 (d, *J* = 8.5 Hz, 2H), 7.16 (dq, *J* = 7.9, 3.1 Hz, 2H), 6.98 (d, *J* = 5.1 Hz, 1H), 6.92 – 6.83 (m, 2H), 5.57 (ddd, *J* = 11.8, 7.3, 4.7 Hz, 1H), 4.68 (t, *J* = 6.9 Hz, 1H), 4.57 (s, 2H), 4.12 (dd, *J* = 5.7, 4.0 Hz, 2H), 3.90 – 3.84 (m, 2H), 3.78 – 3.66 (m, 10H), 3.65 – 3.58 (m, 2H), 3.54 – 3.50 (m, 2H), 3.44 – 3.37 (m, 1H), 3.20 (dd, *J* = 17.8, 4.7 Hz, 1H), 2.68 (d, *J* = 1.5 Hz, 3H), 2.42 (s, 3H), 1.69 (s, 3H).
**$^{13}$C NMR (151 MHz, $CDCl_3$)** δ 170.42, 163.77, 163.67, 158.40, 155.55, 154.30, 149.69, 145.16, 136.59, 136.57, 132.86, 132.06, 131.94, 131.71, 130.78, 130.62, 130.34, 130.20, 129.85, 129.75, 128.56, 128.11, 126.94, 125.02, 114.94, 70.72, 70.51, 70.48, 70.25, 69.66, 69.55, 67.37, 60.08, 54.27, 54.25, 42.05, 41.89, 39.31, 38.98, 29.58, 14.29, 12.97, 11.71.
**HRMS (+ESI):** calcd. $C_{44}H_{47}BrCl_2N_7O_6S$ for = 950.1863; found 950.1852

**Supplementary Table Legends**

**Supplementary Table 1. Structures of covalent ligands screened against RNF114**

**Supplementary Table 2. IsoTOP-ABPP analysis of EN219 in 231MFP breast cancer cells.** IsoTOP-ABPP analysis of EN219 in 231MFP breast cancer cells. 231MFP cells were treated *in situ* with DMSO vehicle or EN219 (1 μM) for 90 min. Control and treated cell lysates were labeled with IA-alkyne (100 μM) for 1 h, after which isotopically light (control) or heavy (EN219-treated) biotin-azide bearing a TEV tag was appended by CuAAC. Proteomes were mixed in a 1:1 ratio, probe-labeled proteins were enriched with avidin and digested with trypsin, and probe-modified peptides were eluted by TEV protease and analyzed by LC-MS/MS.

**Supplementary Table 3. TMT-based proteomic analysis of EN219-alkyne pulldown competed by EN219.** 231MFP cells were treated with DMSO vehicle or EN219 (20 μM) 30 min prior to treating cells with DMSO or EN219-alkyne probe (2 μM) for 90 min. Resulting cell lysates were subjected to CuAAC with biotin-azide to append a biotin enrichment handle onto EN219-alkyne labeled proteins *ex situ*. EN219-alkyne labeled proteins were subsequently avidin-enriched, digested with trypsin, and resulting tryptic peptides from each treatment group were labeled with TMT reagents and combined and fractionated for LC-MS/MS analysis.

**Supplementary Table 4. TMT-based proteomic analysis of ML 2-14-mediated protein level changes in 231MFP cells.** TMT-based quantitative proteomic data showing ML 2-14-mediated protein level changes in 231MFP cells. 231MFP cells were treated with DMSO vehicle or ML 2-14 for 8 h.

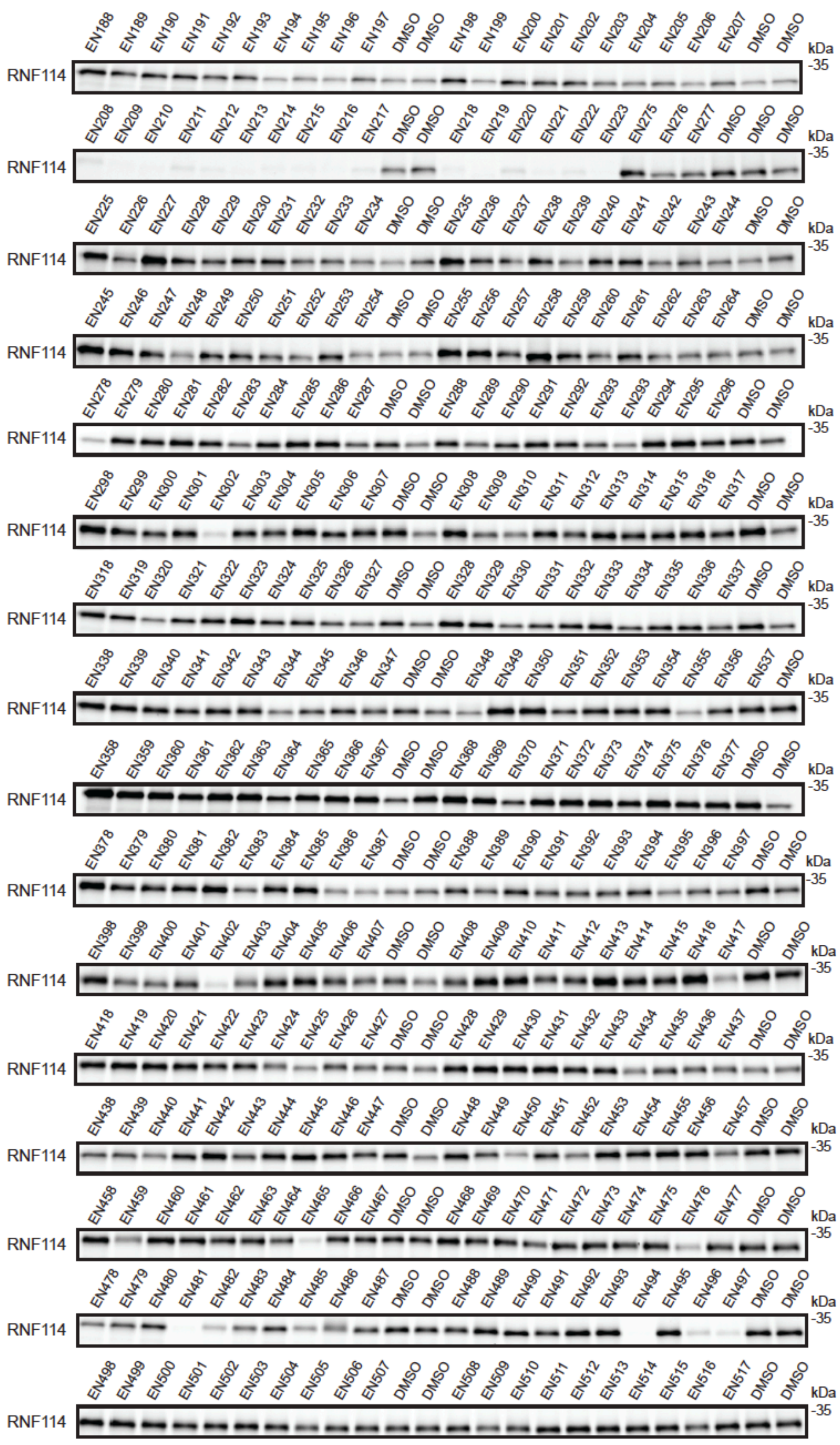

**Supplementary Fig. 1.** Gel-based ABPP assay for screening covalent ligands against IA-rhodamine probe binding to pure RNF114 protein. Loss of fluorescence indicates covalent ligand binding to a cysteine on RNF114. DMSO vehicle or covalent ligands (50 μM) were pre-incubated with pure RNF114 protein (0.1 μg) for 30 min prior to addition of IA-rhodamine (100 nM) for 30 min at room temperature. Proteins were separated by SDS/PAGE and in-gel fluorescence was quantified. The structures of all the compounds screened can be found in **Supplementary Table 1.** This figure is related to **Figure 2.**

a

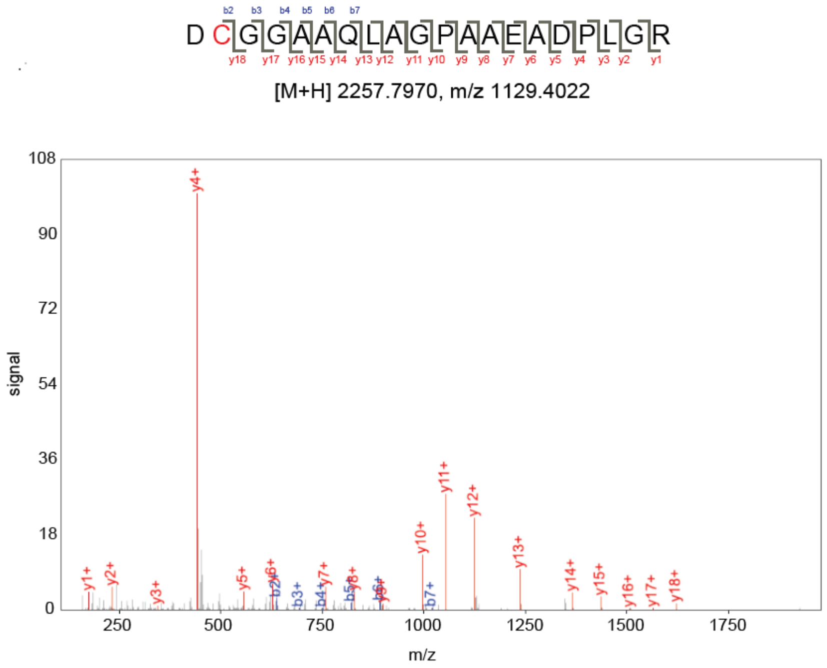


b

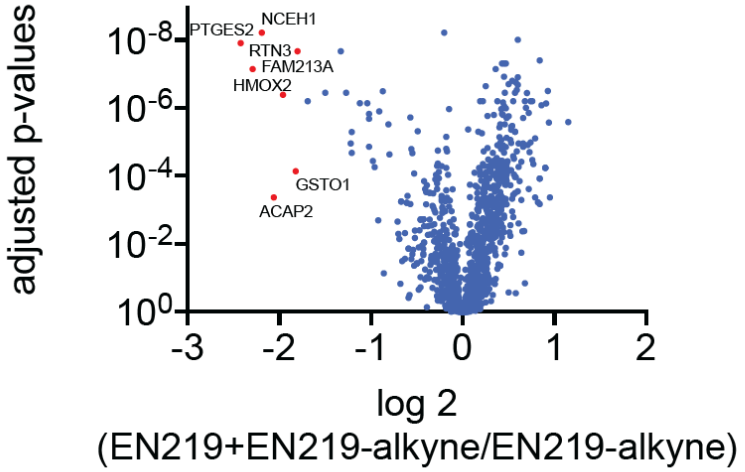


**Supplementary Fig. 2. EN219 reactivity with RNF114 pure protein and TMT-based quantitative proteomic profiling of EN219-alkyne-enriched targets in 231MFP cells. (a)** Pure RNF114 protein was incubated with EN219 (50 μM) for 1 h and subsequent tryptic digests of RNF114 protein were analyzed by LC-MS/MS to look for EN219 covalent modification. The Cysteine 8 highlighted in red was found to be modified. Shown is the mass spectra of the EN219 covalent adduct on the RNF114 tryptic peptide. **(b)** 231MFP cells were treated with DMSO vehicle or EN219 (20 μM) 30 min prior to treating cells with DMSO or EN219-alkyne probe (2 μM) for 90 min. Resulting cell lysates were subjected to CuAAC with biotin-azide to append a biotin enrichment handle onto EN219-alkyne labeled proteins *ex situ*. EN219-alkyne labeled proteins were subsequently avidin-enriched, digested with trypsin, and resulting tryptic peptides from each treatment group were labeled with TMT reagents and combined and fractionated for LC-MS/MS analysis. Shown are average TMT ratios and adjusted p-values comparing EN219 pre-treated EN219-alkyne labeled groups to EN219-alkyne labeled groups to identify EN219-alkyne-labeled proteins that are competed by the parent compound EN219. Shown in red are proteins that show EN219+EN219-alkyne versus EN219-alkyne ratios <0.33 with adjusted p<0.05. The full dataset can be found in **Supplementary Table 3.** This figure is related to **Figure 2.**

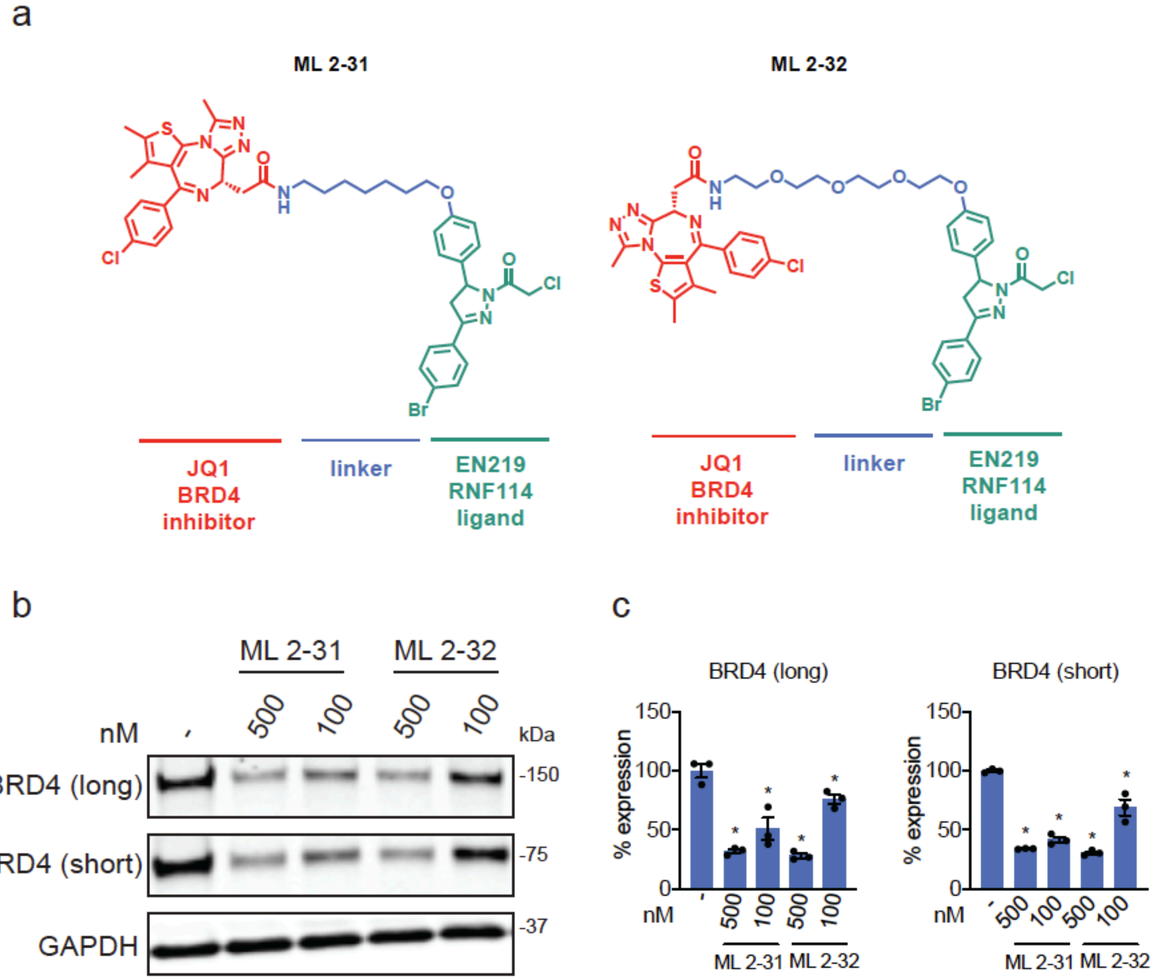


**Supplementary Fig. 3. EN219-based BRD4 degraders. (a)** Structure of ML 2-31 and ML 2-32, EN219-based BRD4 degraders linking EN219 to BET inhibitor JQ1 with two different linkers. **(b)** Degradation of BRD4 by ML 2-31 and ML 2-32. 231MFP cells were treated with DMSO vehicle or ML 2-31 or ML 2-32 for 8 h and the long and short isoforms of BRD4 and loading control GAPDH were detected by Western blotting. **(c)** Percentage of BRD4 degradation quantified from **(b)**. Data shown in **(c)** are average and individual replicate values. Blots shown in **(b)** are representative of n=3 biological replicates/group. This figure is related to **Figure 3.**

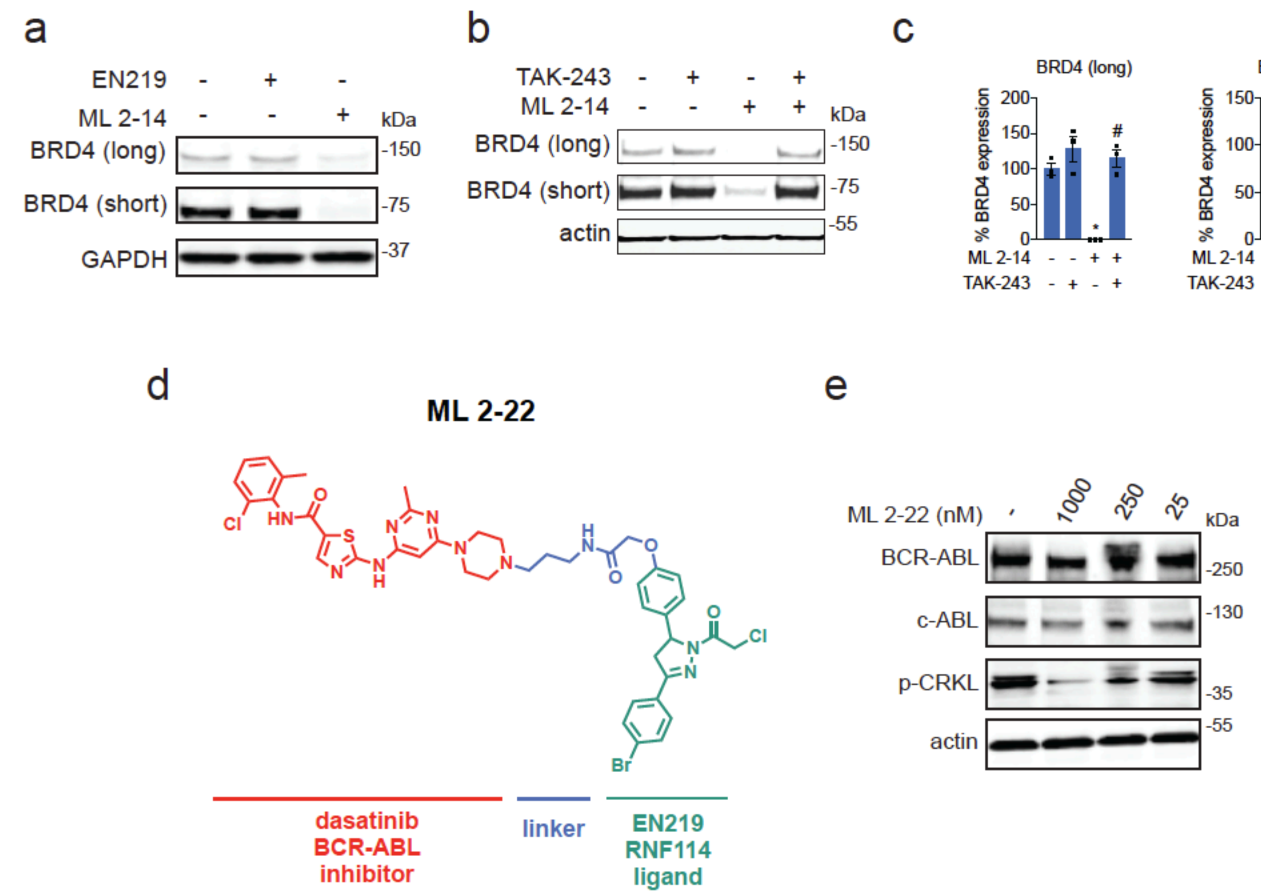


**Supplementary Fig. 4. ML 2-14 and ML 2-22 mediated degradation of BRD4 and BCR-ABL, respectively.** **(a)** BRD4 and loading control GAPDH expression in 231MFP cells treated with DMSO, EN219 (100 nM), and ML 2-14 (100 nM) for 20 h detected by Western blotting. **(b)** Ubiquitin activating enzyme (E1)-dependent degradation of BRD4 by ML 2-14. 231MFP cells were treated with DMSO vehicle or with E1 inhibitor TAK-243 (10 μM) 30 min prior to DMSO vehicle or ML 2-14 (100 nM) treatment for 8 h. BRD4 and loading control actin levels were detected by Western blotting. **(c)** Quantification of data from **(b).** **(d)** Structure of ML 2-22 degrader that linked EN219 to dasatinib. **(e)** BCR-ABL, c-ABL, p-CRKL, and loading control actin levels assessed by Western blot from K562 cells treated with DMSO vehicle or ML 2-22 for 20 h. Blots in **(a, b, e)** are a representative from n=3 biological replicates/group. This figure is related to **Figure 3** and **Figure 4.**
